# Molecular Alterations of Bovine Serum Albumin Induced by the Food Dye Acid Yellow 23: A Mechanistic Study

**DOI:** 10.64898/2026.07.08.737154

**Authors:** Parshant Dahiya, Alpana Verma, Vishal Mevada, Satish Kumar, Neelkant Verma

## Abstract

The widespread use of synthetic food dyes, such as Acid Yellow 23 (AY 23), in the food, cosmetics, and pharmaceutical industries raises questions about their potential effects on biological systems and public health. The concentration-dependent interaction between AY 23 and bovine serum albumin (BSA), a crucial model protein for understanding pharmacokinetics and protein-ligand behaviour, was examined in this study. We demonstrate that, under physiological conditions, increasing the dye concentration from 50 µM to 200 µM leads to notable conformational changes, increased surface hydrophobicity, and protein aggregation, as determined using a multimodal biophysical approach including fluorescence spectroscopy. Direct visualisation verified these structural changes and aggregate formation, whereas hemolytic assay confirmed the high hemolytic nature of AY 23-induced fibrils. Additionally, this study provides a mechanistic basis for the toxicological effects of AY 23, underscoring the implications of food dyes for public health.

**Highlights:**

- Acid Yellow 23 (AY 23) modulates Bovine Serum Albumin (BSA) structure and leads to aggregation under physiological conditions.
- Structural alteration is followed by binding of AY 23 at the hydrophobic regions of BSA, causing perturbation of the protein.
- Confirmational changes induced by AY 23 in the BSA via interaction with Asp108, Pro110 and Ala193.
- Protein intermediates formed after AY 23 interaction are observed to be hemolytic in nature.
- Molecular mechanism of AY 23-mediated perturbation of BSA was assessed.

**Graphical Abstract:** 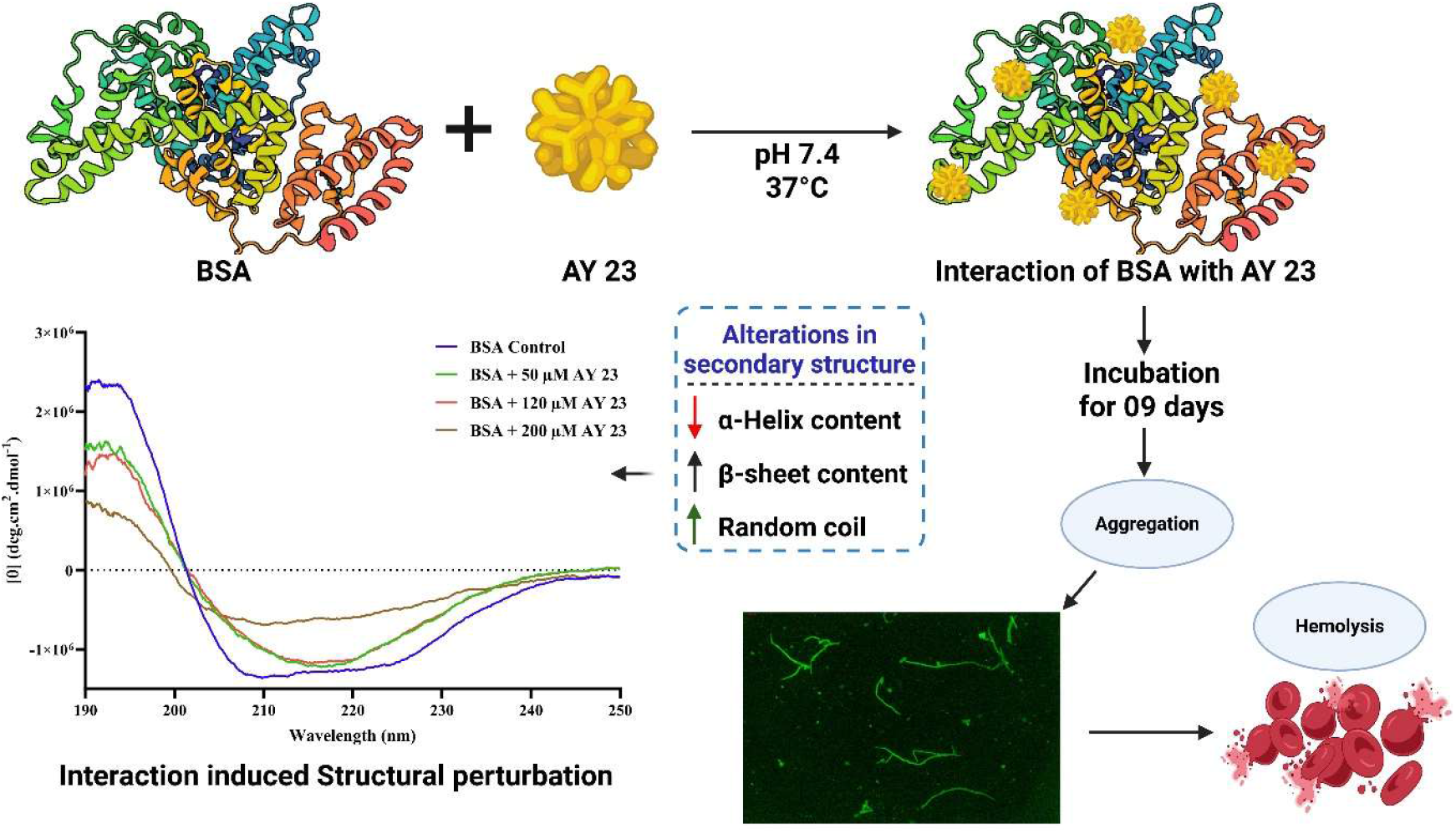

## 1. Introduction

The extensive use of artificial additives in processed foods has raised questions about their safety and potential adverse health effects, especially for young children and other susceptible populations. Dyes have been used for centuries as a food additive to improve the appearance of food (Branen et al., 2001). Artificial food dyes are categorised into azo dyes, triphenylmethane dyes, xanthene dyes and quinophthalone dyes (König, 2015). The azo group, identified by a double bond between two nitrogen atoms (N = N), is typically situated between two aromatic rings and is commonly linked with functional groups like the amino (-NH2) or sulfonic (-SO3H) group (Benkhaya et al., 2020). Acid Yellow 23 (AY 23), like other azo dyes, is an effective colouring substance due to its vivid colour, which is created by an azo chromophore. Despite the obvious industrial benefits of these dyes, academics and regulators continue to debate their safety (Amchova et al., 2015). The lack of agreement regarding their health risks has led to inconsistent regulatory frameworks worldwide. Several reports highlights issues including allergic reactions in sensitive individuals, possible behavioural disturbances like hyperactivity in children, and experimental findings showing potential genotoxic or cytotoxic effects (Keener, 2022; Mota et al., 2023). Evaluating the industrial benefits of synthetic dyes in light of the potential health risks emphasizes the necessity for well-structured studies to thoroughly explore their interactions with biological systems.

AY 23, also known as Tartrazine, FD&C Yellow 5, CI 19140 and E102, is a widely used synthetic azo dye with the chemical formula C_16_H_9_N_4_Na_3_O_9_S_2_ and a molecular weight of 534.38 g/mol. It’s structure contains the characteristic azo linkage (-N=N-) as well as sulfonate groups that contribute to its high solubility in water (Leulescu et al., 2018). Although AY 23 is commonly present in many consumer products, its regulation is not uniform worldwide, with some countries adopting more stringent controls than others. An acceptable daily intake (ADI) of 0-7.5 mg/kg and 0-10 mg/kg body weight per day has been established (Amchova et al., 2024); however, various studies have reported toxic effects at similar dosage including hematotoxicity, reprotoxicity and neurological alterations in animal models (Abd-Elhakim et al., 2018; Axon et al., 2012; El-sakhawy et al., 2019). In a study by pan et al., 2011, tartrazine (AY 23) caused molecular alterations in human serum albumin (HSA) and bovine serum albumin (BSA) (Pan et al., 2011). In another study tartrazine (AY 23) caused the fibrillation of HSA and BSA at different pH conditions (Al-Shabib et al., 2018a). In some recent studies, AY 23 was reported to cause fibrillation of cytochrome c, at pH 7 and alterations in β-lactoglobulin (Liu et al., 2026; Ramesh and Bhuyan, 2024). These studies highlighted the potential of AY 23 to cause molecular alterations and protein misfolding at different pH conditions. However, the role of AY 23 interaction with serum albumin at physiological conditions leading to deleterious effects remained unanswered.

Serum albumin, the main cargo protein in plasma, is essential for many physiological processes, especially the binding, transportation, and systemic administration of metal ions, amino acids, fatty acids, and medications. Bovine serum albumin (BSA), a model protein with molecular weight of ∼66 kDa, 583 amino acids and 17 disulfide bonds, BSA has three domains (I, II, III) and two Trp residues (212, 134) and a distinctive heart shaped structure (Carter and Ho, 1994; Mavani et al., 2023). BSA is a well-established globular protein and frequently used, especially for research on protein-ligand interactions and conformational dynamics.

To gain a basic understanding of the molecular mechanism involved in the interaction and structural changes in this major macromolecule, we examined the concentration-dependent interaction between AY 23 and BSA under physiological conditions at temperature of 37 ℃ and pH 7.4. Our study aimed to provide a mechanistic insight of AY 23 induced structural and confirmational changes of BSA. This study confirms the role of AY 23 in the modulation of BSA leading to toxic fibrillation and underscores the potential deleterious effects of this food dye.

## 2. Materials and Methods

### 2.1. Chemicals and Reagents

Bovine Serum Albumin (BSA Fraction V >98% purity, CAS #9048-46-8) was purchased from Hi-Media (India). Acid Yellow 23 (purity >98%, CAS #1934-21-0) (CI 19140) was obtained from TCI Chemicals (India), Thioflavin T, Nile Red, sodium phosphate monobasic, sodium dodecyl sulfate (SDS), acrylamide, N, N’-methylene bisacrylamide (Bis), N, N, N’, N’-tetramethylethylenediamine (TEMED), and ammonium persulfate (APS) were obtained from Hi-Media (India). All other chemicals used were of analytical grade.

### 2.2. Sample Preparation

The 20 mM sodium phosphate buffer solution was prepared, filtered through a 0.22 μM filter and adjusted to pH 7.4 using a Thermo Scientific Orion Star A211 digital pH meter at room temperature. Stock solutions of Bovine Serum Albumin (BSA) were prepared in double-deionised water, and the protein concentration was accurately quantified by measuring its absorbance at 280 nm, utilising a molar extinction coefficient of 43,824 M⁻¹cm⁻¹ (Ansari et al., 2025). A working concentration of 30 μM of BSA was used for all experiments. AY 23 was dissolved in double-deionised water to a 1 mM stock solution. The binding studies used a constant BSA concentration of 30 μM, with AY 23 concentrations of 50 μM, 120 μM, and 200 μM to examine concentration-dependent effects. All samples, including control samples containing only BSA or AY 23 (at varying concentrations), were made in triplicate. The samples were then incubated at 37°C for different time points for measurement.

### 2.3. Molecular Dynamics Simulation

The Crystal Structure of Bovine Serum Albumin (BSA) as downloaded from the Protein Databank (PDB) having the PDB ID 4F5S. While the ligand included in this study, Acid Yellow 23 (AY 23), was retrieved from PubChem with Chemical ID:16013. Both these structures were pre-processed by removing water molecules and adding polar hydrogen using the prepare_ligand.py and prepare_receptor.py scripts available in the PyRx v.1.0 software. The grid was defined for molecular docking, and Docking was performed using AutoDock Vina (v1.0) with an exhaustiveness of 20. The grid center was kept to x = 9.81208101552, y=21.6569, z = 102.441307998 and size to x = 113.017195148, y = 64.6051993847, z = 85.0043848422. The docking results were visualised using Pymol 3.0. The best pose was selected based on the lowest binding energy for the protein-ligand complex. The resulting complex was further analysed using BIOVIA Discovery Studio Visualizer to identify the interacting amino acids.

The molecular dynamics simulations were performed using the “Desmond V package” (Schrodinger, 2024) to investigate changes in the protein-ligand complex formed between BSA (PDB ID: 4F5S) and the ligand (CID: 16013). For the ligand molecule, the OPLS force field was used for the complex (Al-Shabib et al., 2018b). An orthorhombic cubic box was utilised for molecular dynamics simulations, with the protein-ligand complex centred within it. TIP3P water molecules filled the environment, maintaining a minimum distance of 10 Å from the box edge to the protein atoms. The boundary condition box volume was calculated based on the complex type, and to neutralise the system, Na+ and Cl-counter ions were randomly distributed throughout the box. This setup accurately reflects the biological context for dynamical studies. The energy minimisation was performed using the steepest descent (SD) method with up to 2,000 iterations, and the conjugate-gradient (CG) algorithm with 1,000 iterations and a convergence threshold of 50e. The production run for molecular simulation was performed for 100ns. The system was slowly heated to maintain a temperature of 300 K and a pressure of 1 bar using the Nose-Hoover thermostatic algorithm and the Martina-Tobias-Klein method. The Particle Mesh Ewald (PME) method was utilised to compute long-range electrostatic interactions, with a grid spacing of 0.8 Å. The results were examined by looking at the RMSD (root mean square deviation) values for both the protein and the ligand, as well as the fluctuations in their positions (RMSF).

### 2.4. UV-Visible Spectroscopy

UV-visible spectroscopy was performed to confirm the interaction of AY 23 and BSA. A Lab India dual-beam UV-visible spectrophotometer (Next Gen Series) was used to confirm the interaction between AY 23 and BSA. The baseline correction was made using sodium phosphate buffer. The final concentration of the BSA was kept at 30 µM. Absorption spectra of BSA with and without varying concentrations of AY 23 (10 µM – 210 µM; with 30 µM step up) were recorded from 200 nm to 600 nm at 298 K, thrice for each sample.

### 2.5. Intrinsic fluorescence measurements and Stern–Volmer analysis

Bovine serum albumin (BSA) and Acid Yellow 23 (AY 23; tartrazine) were used to investigate the interaction between the food colorant and the protein. BSA concentration was maintained at 30 µM throughout the fluorescence measurements. Steady-state fluorescence measurements were performed at 298 K using a Jobin–Yvon Fluoromax-4 spectrofluorometer (Horiba Scientific) equipped with a xenon lamp and a 1-cm quartz cuvette. The intrinsic fluorescence of BSA was monitored using an excitation wavelength of 295 nm, and emission spectra were recorded over 300 to 450 nm. The instrumental excitation and emission slit widths were 1.5 nm and 10 nm, with a scan rate of 1 nm min⁻¹. BSA was analyzed in the absence and presence of increasing concentrations of AY23 (0, 10, 30, 60, 90, 120, 150, 180 and 210 µM). The final result was obtained by subtracting blanks from the average of five distinct spectra.

### 2.6. Circular Dichroism (CD) Spectroscopy

Circular Dichroism (CD) measurements were performed on a JASCO J-815 spectropolarimeter to analyse the secondary structure of 30 µM BSA incubated with varying concentrations of AY 23 (50, 120, and 200 µM). A 0.1 cm path-length quartz cuvette was used, with all experiments conducted at 25°C. CD spectra were acquired in the far-UV region (190-250 nm) by averaging 20 consecutive scans for each sample. The scan rate was set to 50 nm/min, with a response time of 1 second and a bandwidth of 1 nm. The spectrum of each sample was subtracted from the blank. The BeStSel was used to process and analyse all spectral data (Micsonai et al., 2025).

### 2.7. Dynamic Light Scattering (DLS)

DLS measurements were used to track changes in the hydrodynamic size of BSA and assess the formation of aggregates. BSA samples with various concentrations of AY 23 (50, 120, and 200 µM) were put in a quartz cuvette. The samples were analysed using a Zetasizer Nano S90 DLS (Malvern Instruments Ltd, UK) at 25°C (Varma et al., 2018). The hydrodynamic radii (Rh) of each sample were calculated using Malvern Zetasizer Software.

### 2.8. Turbidity

Changes in sample turbidity over time were monitored to track the development and expansion of protein aggregates. A UV-visible spectrophotometer was used to assess turbidity at 550 nm, where neither AY 23 nor BSA shows any discernible intrinsic absorbance, ensuring that the signal mainly reflects light scattered by large particles. The absorbance at 550 nm was measured after incubating BSA samples (30 μM) with different doses of AY 23 (50 μM, 120 μM, and 200 μM) at 37°C. The size and quantity of the protein aggregates increase as the absorbance at this wavelength increases, which is exactly proportional to the amount of light scattered.

### 2.9. Sodium Dodecyl Sulphate Polyacrylamide Gel Electrophoresis (SDS-PAGE) Assay

SDS-PAGE was utilised to observe the changes in BSA structure in the presence of AY 23. BSA was incubated with AY 23 at various concentrations (50, 120, and 200 µM) and at day 09, samples were subjected to a 12% gel along with a protein ladder. Electrophoresis was carried out with a BIO-RAD Mini-PROTEAN® Tetra System (BIO-RAD, USA) device at a constant voltage of 120V. After electrophoresis, the gels were stained with 0.025% (w/v) Coomassie Brilliant Blue and then subjected to multiple destaining steps. Protein bands were visualised using a Bio-Rad ChemiDoc MP Imaging System.

### 2.10. Nile Red (NR) Binding Assay

To monitor the hydrophobic exposure of BSA protein after the interaction with AY23, Nile Red binding assay was employed. Nile Red, a benzophenoxazone dye, exhibits a strong increase in fluorescence upon binding to a hydrophobic environment (Sackett and Wolff, 1987). Nile red stock solution was prepared by dissolving it in DMSO, and the concentration was determined using the molar extinction coefficient of 19,600 M^-1^ cm^-1^ at 552 nm (Castro et al., 2005). Prior to use, the stock solution was filtered using a 0.22 μM filter and further diluted to 5 μM (<0.5% DMSO) with preincubated BSA samples with and without AY 23. Further the samples subjected to incubation for 20 minutes in dark and subsequently excited at 550 nm and emission spectra recorded from 565 to 750 nm using a Jobin–Yvon Fluoromax-4 spectrofluorometer (Horiba Scientific). The excitation and emission slit width set at 5 nm each and obtained spectra subtracted by their respective blanks.

### 2.11. Thioflavin T Fluorescence Microscopy

Fluorescence microscopy was utilised to directly visualise BSA aggregation induced by AY 23. Thioflavin T (ThT) fluorescent probe is used to stain protein aggregates at a protein-to-dye molar ratio of 1:2 for 30 minutes in the dark. A ten-microlitre sample of BSA incubated with varying concentrations of AY 23 (at day 09) was prepared on glass slides (Demeule et al., 2007). Images were acquired using a Nikon DM 5200 Fluorescence microscope with 40x and 100x oil-immersion objectives. Morphological changes, aggregate formation, and spatial distribution of the dye-protein complexes were analysed.

### 2.12. Atomic Force Microscopy (AFM)

To examine surface topography and the structures of BSA and its complexes, incubated samples (on day 09) with or without AY 23 were deposited onto freshly cleaved muscovite mica substrates. After drying the sample-loaded substrates in a dust-free environment under a 40W lamp for 30 minutes, followed by high-vacuum drying, they were imaged using an atomic force microscope (AFM) (INNOVA, Bruker). Imaging was performed in acoustic AC (tapping) mode at room temperature in air. A silicon nitride tip (NSC 12(c), MikroMasch) was used with a force constant of 2.0 N/m and a resonant frequency of ∼276 kHz. The images were acquired at a scan speed of 1.5 lines/sec and analysed with Nanoscope Analysis software.

### 2.13. Hemolytic Assay

The hemolytic assay was conducted following prior approval from the Institutional Ethics Committee, National Forensic Sciences University (NFSU), under ethical clearance certificate number NFSU/SDSR/IEC/Certificate/1058/2024. Fresh human blood was obtained from a healthy volunteer after informed consent and used strictly in accordance with the approved ethical guidelines. The collected blood sample was centrifuged at 2000 rpm for 10 min to separate erythrocytes, and the resulting red blood cell (RBC) pellet was washed three times with isotonic phosphate-buffered saline (PBS, pH 7.4). A standardised RBC suspension was then prepared in fresh PBS. The AY 23 and BSA complexes at varying concentrations (on day 09) were incubated with the RBC suspension at 37 °C for 2 hours to determine hemolytic activity. After incubation, samples were centrifuged at 1500 rpm for 10 min, and the absorbance of the supernatant was measured at 405 nm to determine haemoglobin release (Sæbø et al., 2023). Hemolysis was determined relative to erythrocytes treated with deionised water, which served as the positive control (100% lysis), and to RBCs incubated with PBS alone, which served as the negative control (0% lysis). All tests were carried out in triplicate, and the results were reported as mean values. The percentage of lysis was calculated using the following formula:

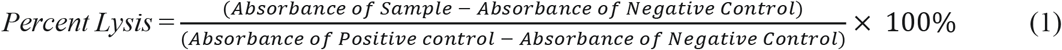

## 3. Results and discussion

### 3.1. Molecular Docking and Molecular Simulation

Molecular docking was performed to investigate the potential binding mode of AY 23 within BSA and to identify amino acid residues potentially involved in stabilizing the protein–ligand complex. AutoDock Vina generated 20 docking conformations with predicted docking scores ranging from −9.7 to −7.46 (ΔGb, kcal/Mol). The top-ranked docking pose (-9.7 kcal/Mol) was further processed for interaction analysis.

The molecular docking analysis revealed that the ligand exhibited binding within the target protein’s binding pocket/binding cavity via multiple non-covalent interactions (Fig. 1). The docking pose demonstrated that the ligand is well accommodated within the binding cavity and forms several hydrogen bonds, electrostatic interactions, and hydrophobic contacts with key amino acid residues (Roy et al., 2019). Specifically, conventional hydrogen bonds were observed between the ligand and residues Asp111, Leu112, and Arg435, which may contribute significantly to binding stabilisation. The interaction with Asp111 occurs through the carboxyl functional group of the ligand, while Leu112 participates in hydrogen bonding with the hydroxyl moiety. Another hydrogen bond is formed between the ligand’s sulfonyl oxygen and Arg435, further anchoring the ligand within the binding site. In addition to hydrogen bonding, electrostatic interactions were identified involving residues His145, Arg458, and Glu424, which participate in π–anion and attractive charge interactions with the aromatic core of the ligand. These electrostatic interactions are likely to enhance the ligand–protein binding affinity (Cao et al., 2024). The ligand also displayed interaction with Asp108 and π–alkyl interactions with Pro110 and Ala193, indicating favourable hydrophobic contacts within the binding pocket. Furthermore, several residues including Arg196, Ser192, Ser428, Ile455, Leu454, Thr190, Tyr451, Ser109, and Pro113 contributed van der Waals interactions, which collectively stabilize the ligand conformation within the active site. These results show a favourable predicted binding score for the formation of a BSA-AY 23 complex.

**Fig. 1.**
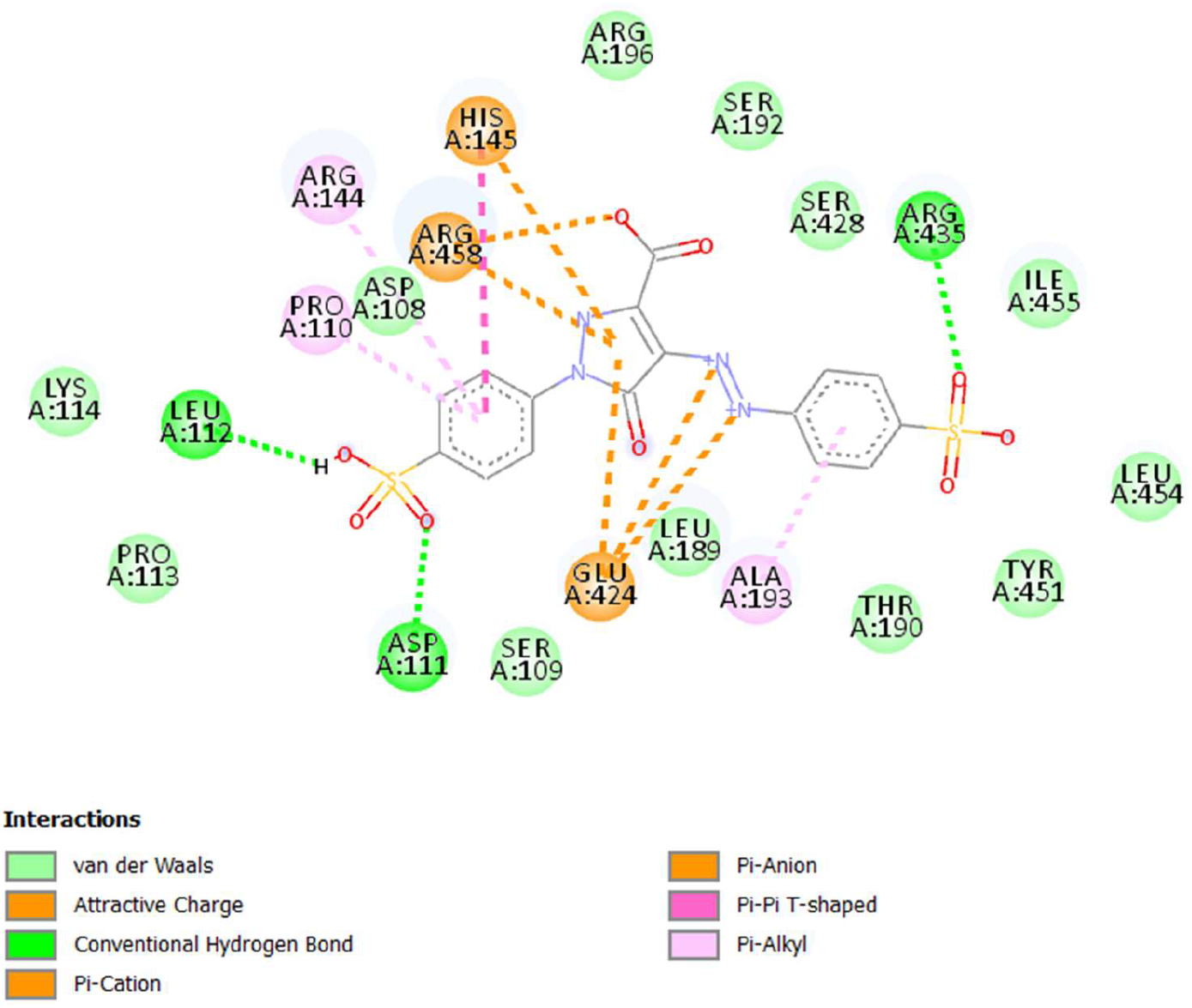
Molecular Docking Reveals Key Amino Acid Residues and Binding Affinity in the AY 23-BSA Complex.

A 100 ns molecular dynamics simulation was carried out under NPT conditions at 300 K to assess the stability of the BSA–AY 23complex. The protein backbone RMSD increased during the initial phase and reached a relatively stable plateau after approximately 20 ns, with fluctuations predominantly between 3.8 and 4.2 Å for the remainder of the simulation (Fig. 2). The RMSD of the ligand relative to the protein backbone (Lig fit Prot) remained within ∼2.5–3.2 Å, suggesting that the ligand stayed within the binding pocket during the simulation. In addition, the internal ligand RMSD (Lig fit Lig) was comparatively lower, fluctuating around ∼1.5–2.0 Å. Collectively, these RMSD profiles indicate that the BSA–AY 23 complex maintained a relatively stable conformational state and that AY 23 remained associated with the predicted binding region throughout the 100 ns simulation.

**Fig. 2.**
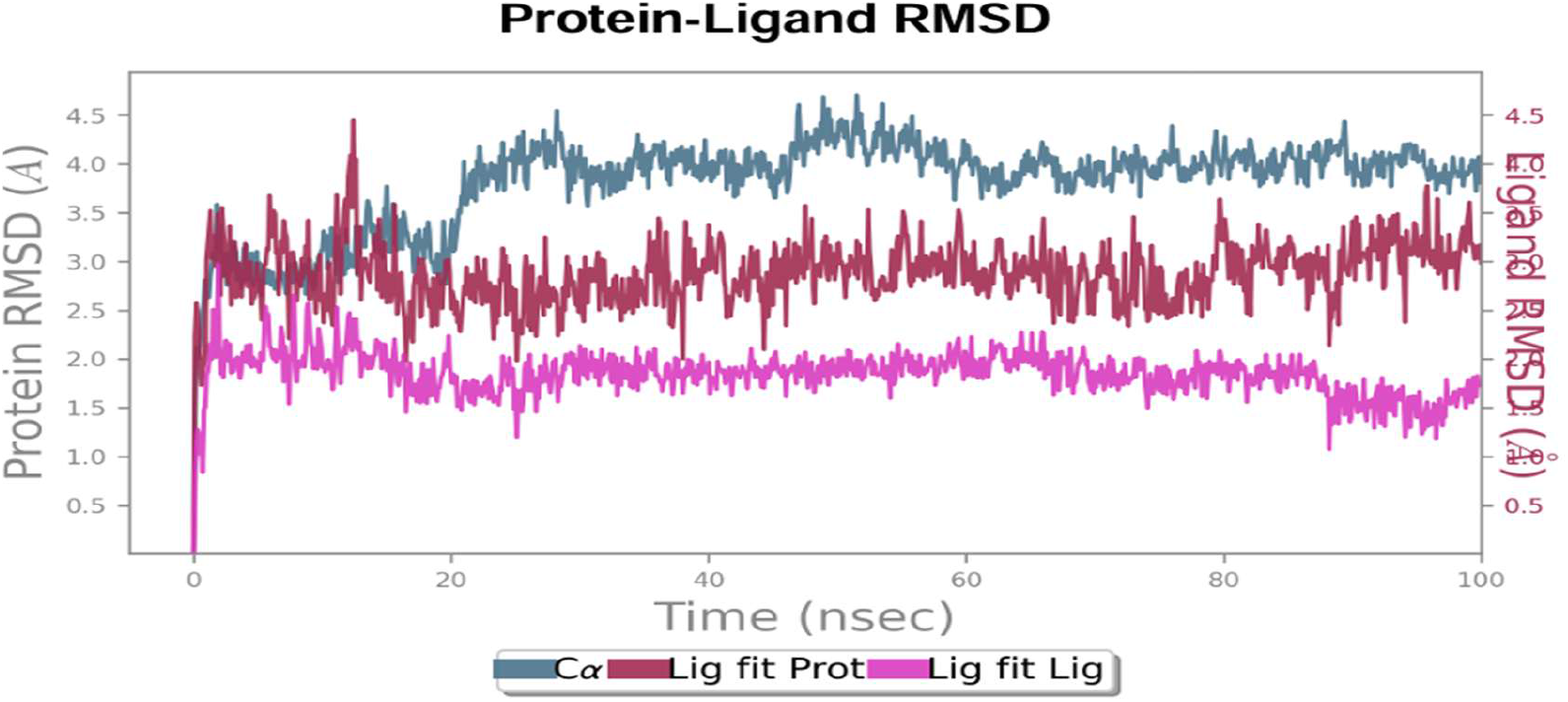
Protein-Ligand Complex RMSD.

The protein’s residue-level flexibility was further evaluated using RMSF analysis (Fig. 3). Most residues exhibited fluctuations between approximately 1.0 and 2.5 Å, whereas higher fluctuations of up to 3.5-4.0 Å were observed mainly in flexible loop regions and terminal segments. Residues within the predicted binding region generally exhibited comparatively low fluctuations during the simulation. Protein-ligand contact analysis revealed hydrogen bonds, hydrophobic interactions, ionic interactions, and water-mediated bridges during the simulation. The persistent interactions with the ligand occurred with several residues, such as LYS114, ARG185, GLU186, THR190, ARG194, SER428, ARG435, TYR451, ARG458, and GLU519 (Fig. 1). The number of protein-ligand contacts generally ranged from approximately 12 to 20 during the trajectory, supporting the continued association of AY 23 with the predicted BSA binding region throughout the simulation.

**Fig. 3.**
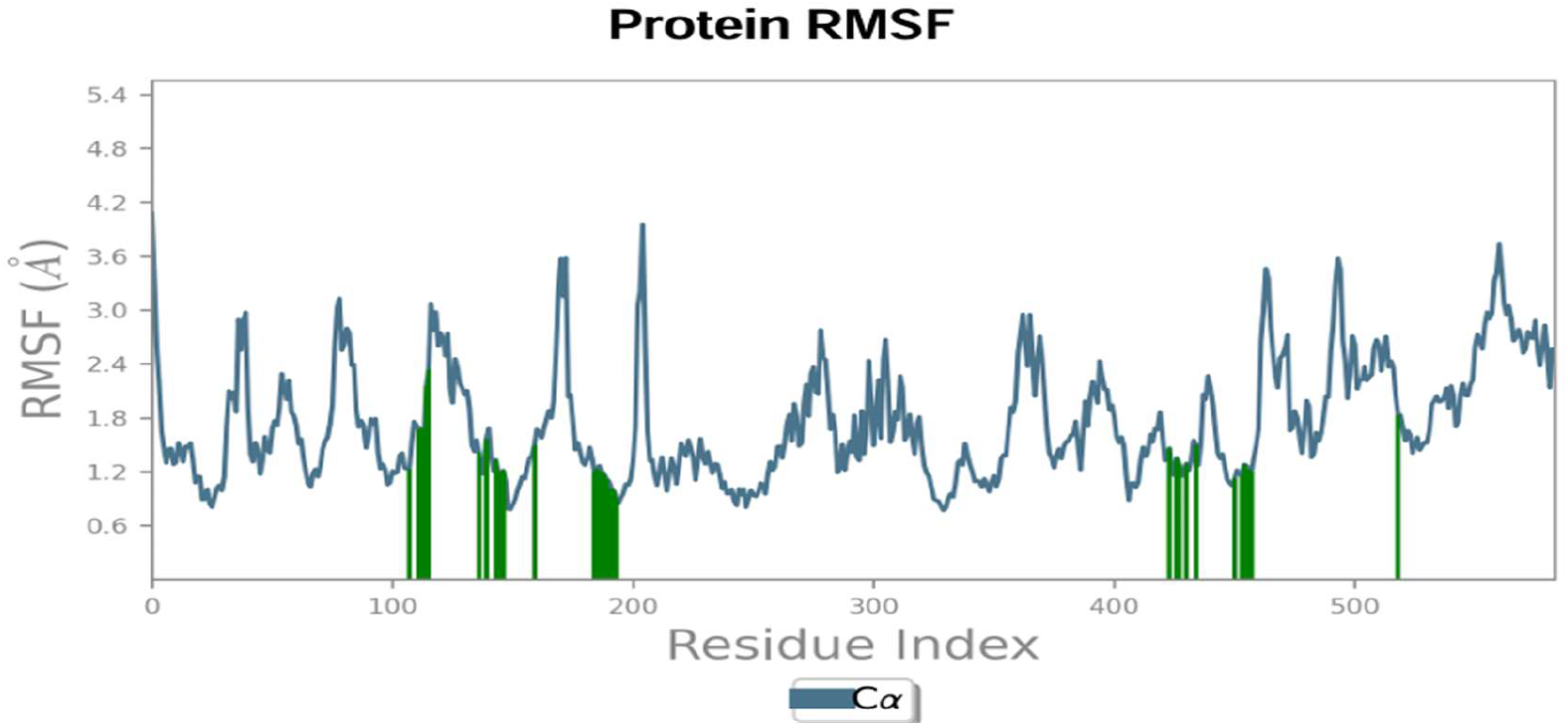
Protein-Ligand Complex RMSF.

### 3.2. UV-Visible Spectroscopy

UV-visible spectroscopy was employed to explore the interaction between AY 23 and bovine serum albumin. Due to the presence of two Trp residues (Trp 134 and Trp 213) in the BSA structure, the conformational changes can be followed using tryptophan at λmax 280 nm. In BSA, the absorption band around 225 nm is primarily attributed to electronic transitions of the peptide backbone, whereas absorption in the 230–300 nm region predominantly arises from the aromatic side chains of Trp, Tyr, and Phe. The absorption around 280 nm is dominated by Trp and Tyr, while Phe exhibits a weaker absorption maximum near 257–258 nm (Pignataro et al., 2020). The UV–visible absorption spectra showed concentration-dependent changes in the spectral profile of BSA following AY 23 exposure (Fig. 4). Native BSA (control) shows the characteristic absorption maxima at ∼280 nm due to aromatic amino acid residues. AY 23 caused a gradual, dose-dependent elevation in absorbance (hyperchromicity) in the 240-320 nm range. The BSA-AY 23 absorption at increasing AY 23 concentrations shifted from 276 nm to 262 nm in a concentration-dependent manner, consistent with a previous study by Y. Liu et al. (Liu et al., 2026). This remarkable blueshift in the BSA absorbance spectra indicates disruption of the tryptophan microenvironment, decreasing the polarity near Trp residues and increasing hydrophobicity. Absorbance patterns of BSA alone (30 µM), AY 23 alone (30 µM) and the BSA-AY 23 complex (1:1) formation were also recorded (Supplementary Figure S1). The difference spectra were calculated and found to be distinct from the absorption spectra of AY 23 alone, suggesting the formation of a ground-state complex between BSA and AY 23 (Dezhampanah et al., 2023).

**Fig. 4.**
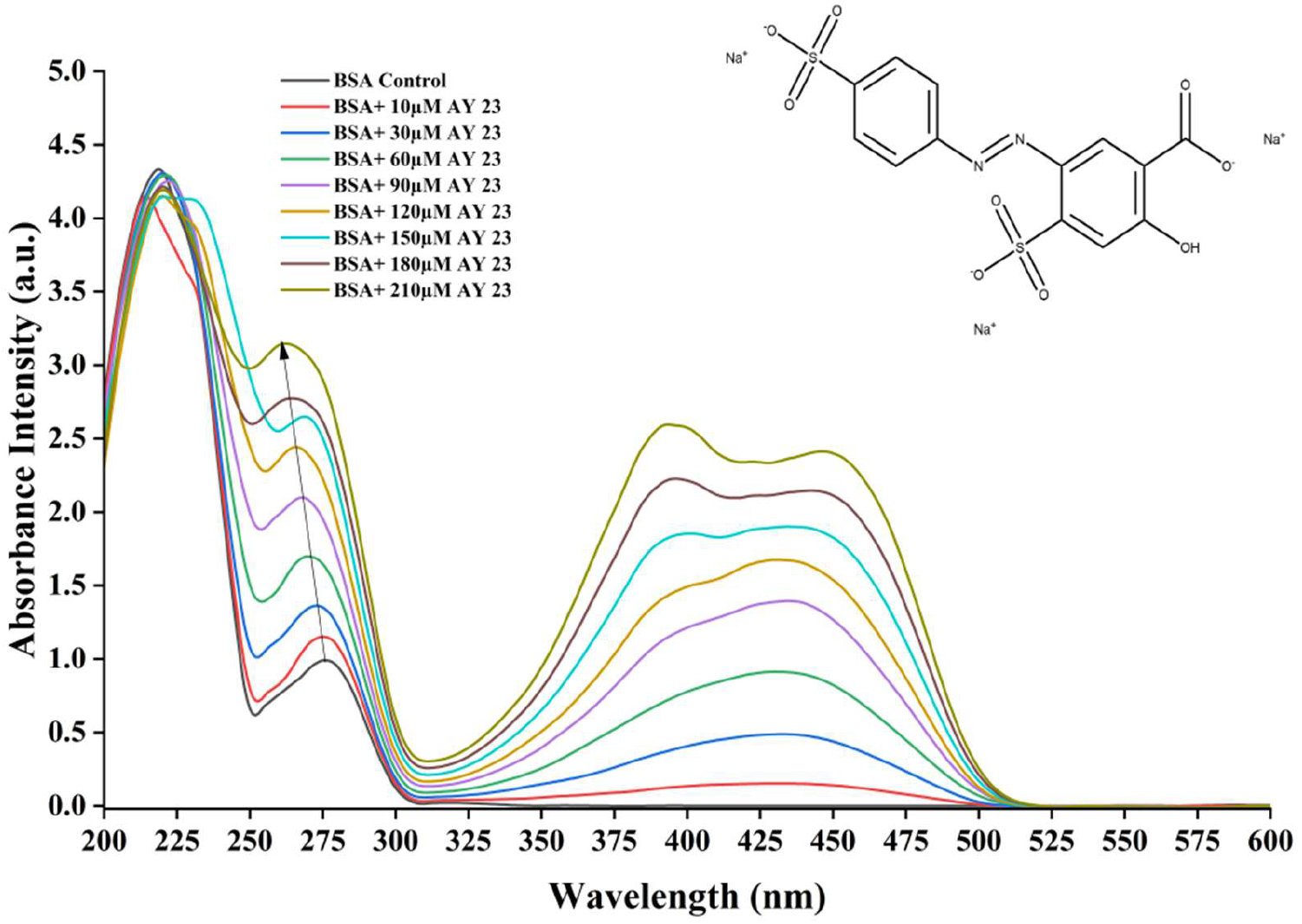
UV-Visible absorption spectra of BSA-AY 23 at different concentrations at pH 7.4.

### 3.3. Stern–Volmer analysis of AY 23-induced fluorescence quenching

The intrinsic fluorescence of BSA arises predominantly from its aromatic residues, particularly tryptophan, making changes in fluorescence intensity sensitive to perturbations in the local environment of these residues. BSA contains two tryptophan residues, Trp-134 and Trp-212, with Trp-212 located within the hydrophobic environment of subdomain IIA and Trp-134 more surface-exposed. Thus, concentration-dependent changes in BSA intrinsic fluorescence were used as an indicator of interaction between AY 23 and the protein. For quantitative analysis, the integrated fluorescence intensity obtained from each emission spectrum was used. The fluorescence intensity in the absence of AY 23 was designated (F_0_), whereas (F) represents the corresponding integrated fluorescence intensity in the presence of AY 23. The Stern–Volmer relationship was expressed as (Lakowicz, 2006):

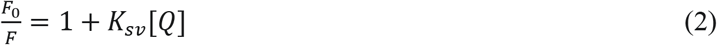

where (F_0_) and (F) represent the fluorescence intensities of BSA in the absence and presence of AY 23, respectively, and [Q] is the AY 23 concentration.

The (F_0_/F) values increased progressively from 1.00 in the absence of AY23 to 9.84 at 210 μM AY 23. The Stern-Volmer plot showed a strong positive linear relationship over the investigated concentration range (10-210 μM), with the regression equation:

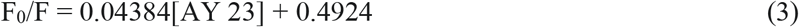

and R² = 0.9858 (Fig. 5). The corresponding slope yielded an apparent Stern-Volmer quenching constant (KSV) of 4.384 × 10⁴ M⁻¹. These results demonstrate a pronounced concentration-dependent attenuation of BSA intrinsic fluorescence following AY 23 exposure. The emission patterns of AY 23 at various concentrations in the absence of BSA are also noted (Supplementary Figure S2), and no contribution of AY 23 to the emission spectra was observed. The fluorescence changes were accompanied by concentration-dependent changes in the UV-Vis absorption profile of BSA. The absorbance at 280 nm increased from 0.9510 for BSA alone to 2.5760 at 210 µM AY 23 (Fig. 4), together with changes in the spectral profile in the aromatic region. The concomitant fluorescence quenching and absorption changes are consistent with an AY 23-BSA ground-state interaction and suggest that static quenching may contribute to the observed fluorescence attenuation, as also reported by previous studies (Liu et al., 2026; Pan et al., 2011).

**Fig. 5.**
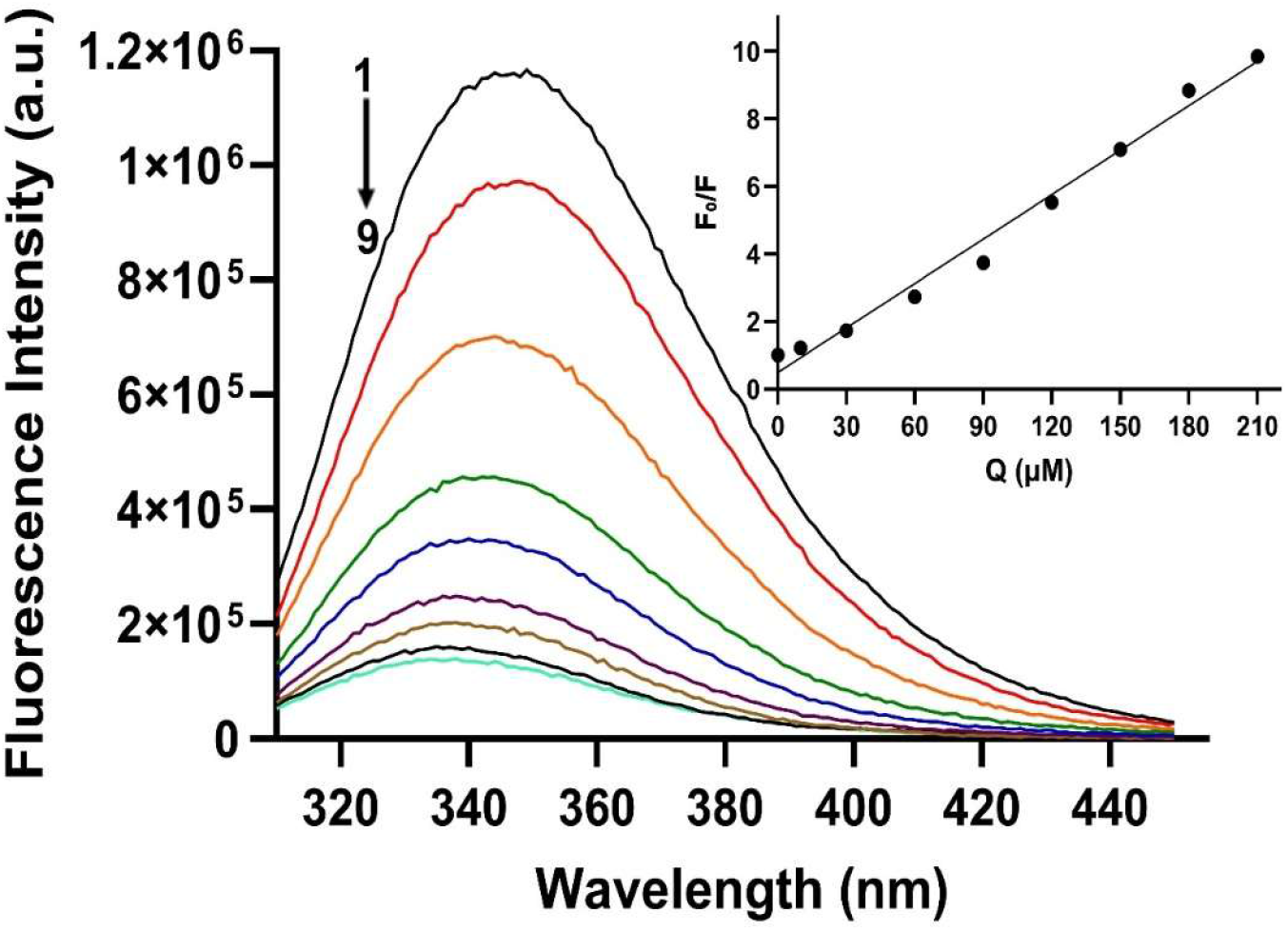
Effect of AY23 concentration on the intrinsic fluorescence of BSA. Fluorescence emission spectra of BSA recorded at 298 K following treatment with increasing concentrations of AY 23. Curves 1–9 correspond to increasing AY 23 concentrations of 0, 10, 30, 60, 90, 120, 150, 180, and 210 μM, respectively. Inset: Stern–Volmer plot of *F*_0_/*F* versus AY 23 (Q) concentration, showing a linear relationship over the investigated concentration range.

### 3.4. Circular Dichroism (CD) Spectroscopy

Circular Dichroism (CD) spectroscopy was performed to investigate the effect of AY 23 on the secondary structure of BSA (Fig. 6). The far-UV CD spectrum of the BSA control exhibited two characteristic negative bands at 208 nm and 222 nm, typical of an α-helical-rich protein. Upon the addition of AY 23, a concentration-dependent decrease in the molar ellipticity was observed, indicating a loss of the native α-helical content of BSA (Waghmare et al., 2018). The spectrum of the 50 μM AY 23 sample showed a slight reduction in α-helical content, while the 120 μM AY 23 sample exhibited a more significant decrease. The most profound structural change occurred at 200 μM AY 23, where the characteristic α-helical bands were almost completely diminished. This confirms that the interaction with AY 23 disrupts BSA’s native folding, leading to a loss of its secondary structure (Koundal et al., 2025). All circular dichroism spectral data were analyzed using BeStSel, which was employed to deconvolute the spectra and estimate the protein secondary structure components such as α-helix, β-sheet, turns, and random coil as shown in Supplementary Table 1.

**Fig. 6.**
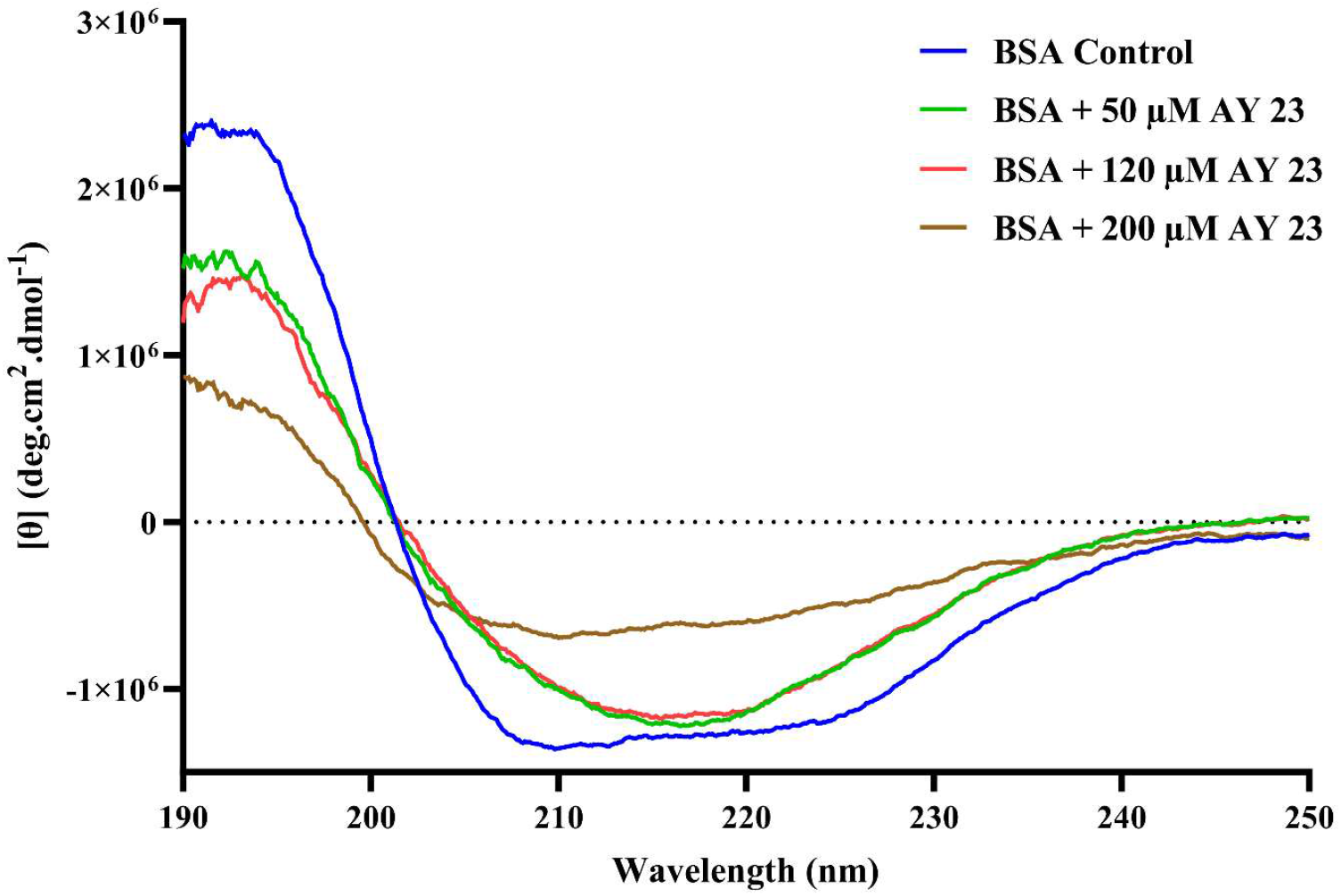
Circular Dichroism (CD) spectroscopy analysis of BSA’s secondary structure changes induced by AY 23.

### 3.5. Dynamic Light Scattering (DLS)

Dynamic Light Scattering (DLS) analysis provides quantitative evidence for the concentration-dependent aggregation of BSA induced by AY 23. DLS was utilized to determine the hydrodynamic radius (Rh) of BSA particles in solution. The hydrodynamic radius (Rh) of the BSA samples was measured, revealing a clear concentration-dependent increase in particle size after 9 days of incubation (Fig. 7). The BSA control showed a Rh of 6.8 ± 1.5 nm, characteristic of the native, monomeric protein (Halder et al., 2021). The addition of AY 23 caused a significant increase in Rh, indicative of protein aggregation. At 50 μM AY 23, the Rh increased to 298.1 ± 112.3 nm, demonstrating that even a low concentration of AY 23 initiates the formation of larger aggregates. Increasing the AY 23 concentrations to 120 μM and 200 μM resulted in a further increase in Rh to 323.7 ± 121.8 nm and 498.6 ± 160.4 nm, respectively. These DLS results provide quantitative evidence that AY 23 promotes BSA aggregation, with the extent of aggregation directly proportional to the dye concentration.

**Fig. 7.**
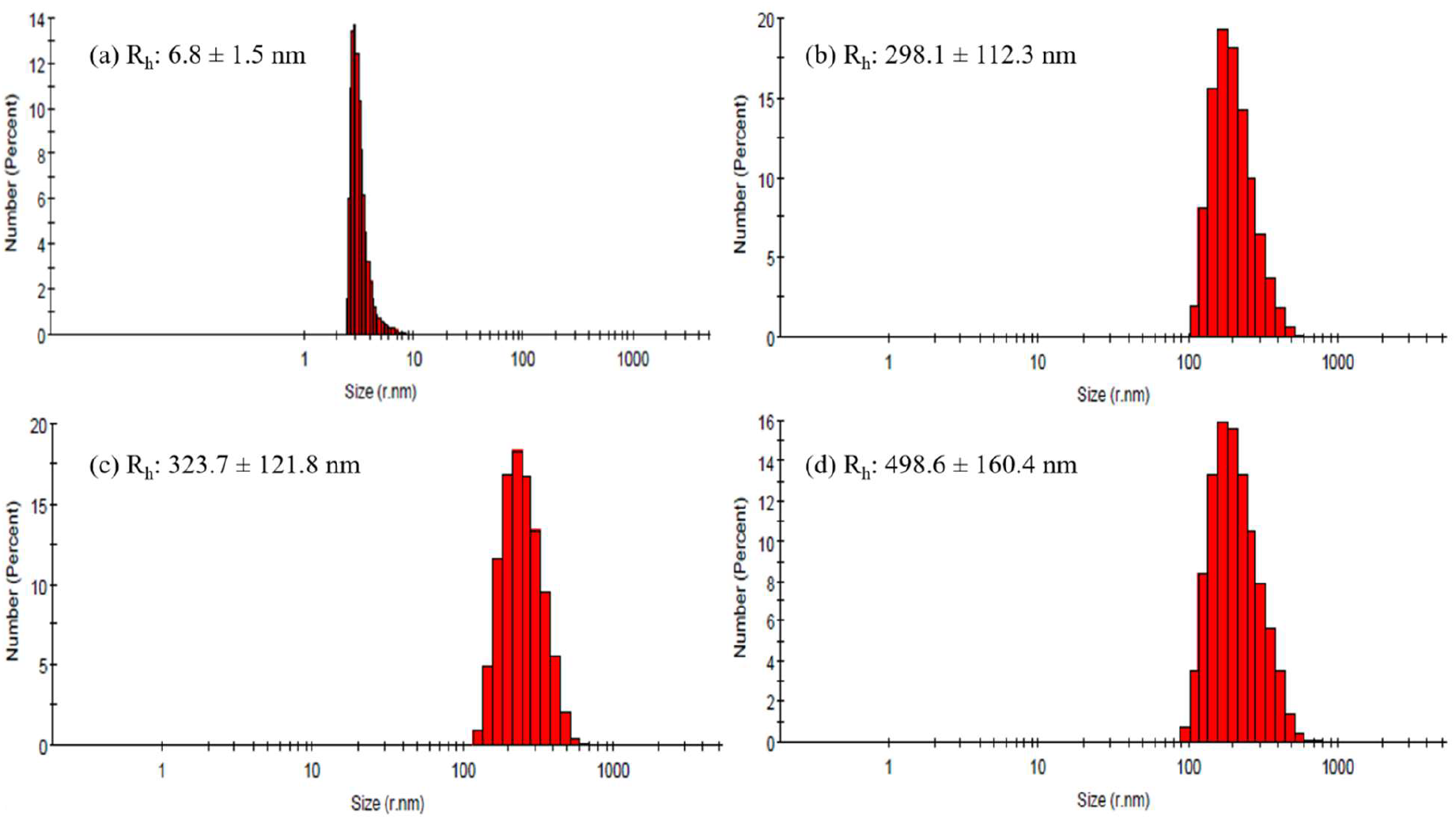
Dynamic light scattering (DLS) analysis showing hydrodynamic radius (Rh) of BSA and its complexes with AY 23 where (a) represents BSA Control; (b) BSA+50 µM AY 23; (c) BSA+120 µM AY 23 and (d) BSA+200 µM AY 23.

### 3.6. Turbidity

To monitor the extent of aggregation turbidity was evaluated at 550 nm (Li et al., 2012). Previous reports suggest that turbidity of aggregated proteins can be studied in the range of 350 – 650 nm (Kawachi et al., 2013). Fig. 8 shows a concentration-dependent increment in turbidity in BSA samples incubated with various concentrations of AY 23. Native BSA had the lowest turbidity, while turbidity increased gradually after adding the dye AY 23, indicating increased aggregation formation. Turbidity increased to 1.25-fold at 50 μM AY 23 compared to native BSA and 1.9-fold at 200 μM at pH 7.4. This steady increase in turbidity indicates that AY 23 promotes BSA aggregation.

**Fig. 8.**
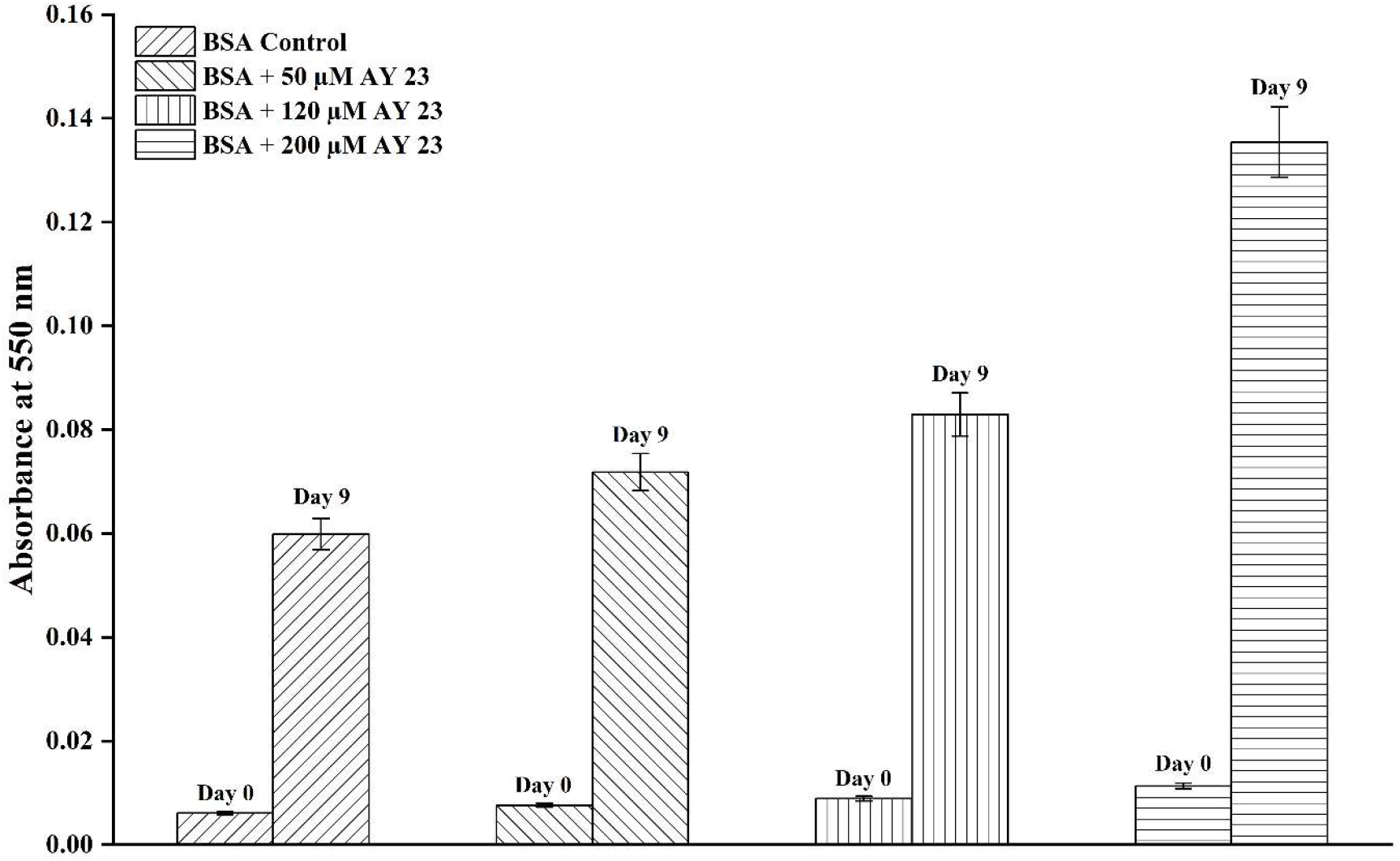
Turbidity assay of bovine serum albumin (BSA) at 550 nm in absence and presence of AY 23 at pH 7.4.

### 3.7. Sodium Dodecyl Sulfate-Polyacrylamide Gel Electrophoresis (SDS-PAGE)

SDS-PAGE was performed under denaturing conditions, with and without AY 23. Fig. 9 shows the results obtained after SDS-PAGE assay; a native band of ∼66 kDa is visible in lane 1; lanes 2 to 4 represent the increasing concentration of AY 23 incubated with BSA at day 9. In Lane 2 and Lane 3, AY 23 shows stacking of higher-order aggregates at the top, and Lane 4 shows a diffusing pattern denoting the presence of aggregation species in the sample (Wang, 2005). This phenomenon indicates that AY 23 can induce BSA monomers to form aggregates.

**Fig. 9.**
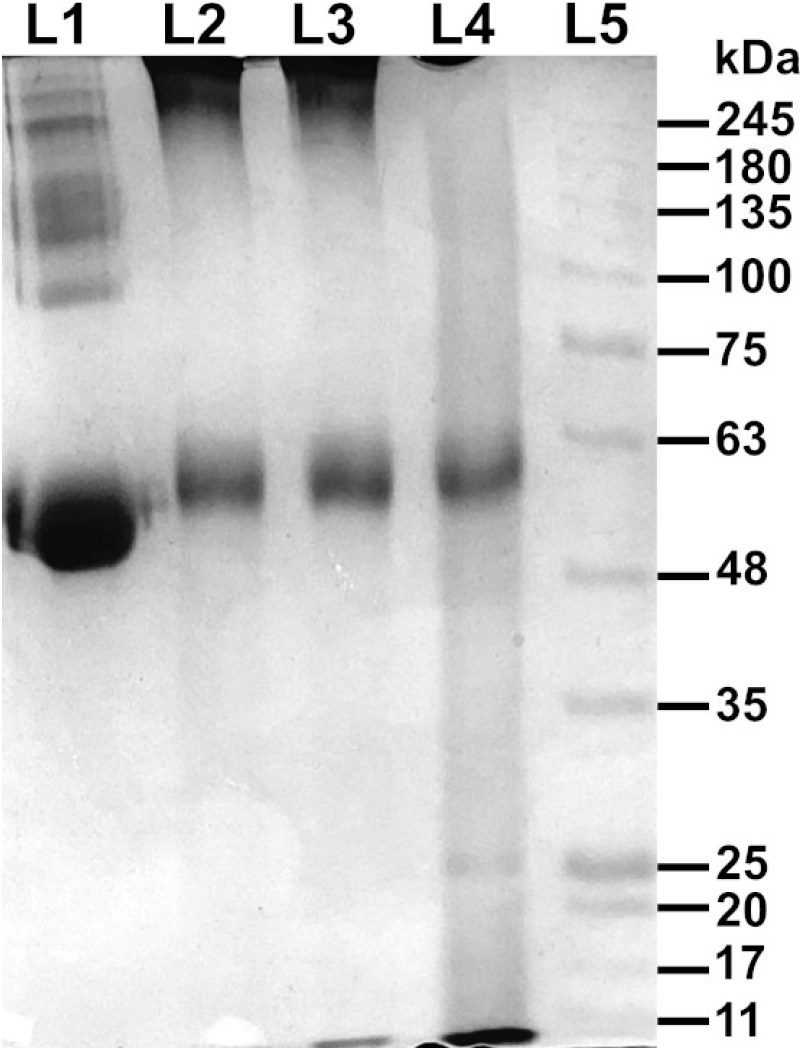
SDS-PAGE analysis of BSA and its interaction with AY 23. Lane 1: BSA (Control); Lane 2: BSA + AY (50 μM); Lane 3: BSA + AY (120 μM); Lane 4: BSA + AY (200 μM); Lane 5: Protein Ladder.

### 3.8. Nile red binding assay

Nile red is a hydrophobic probe as it binds to the solvent-exposed hydrophobic patches during protein unfolding and aggregation (Sackett and Wolff, 1987). A characteristic increase in Nile red fluorescence emission is observed when hydrophobic patches are present. Nile red is a good complementary probe along with ThT assay and DLS, which can reveal the extent of aggregation. In our study, the Nile Red binding assay showed increased fluorescence in BSA samples incubated with AY 23 on day 9 (Fig. 10). This enhancement followed a concentration-dependent pattern as AY 23 concentration increased. This indicates the unfolding of BSA after the interaction with varying concentrations of AY 23, leading to misfolding. It is noted that AY 23 without BSA didn’t exhibit any significant fluorescence, ruling out its contribution. These findings are well supported by the size distribution of BSA samples observed by DLS and further by SDS-PAGE.

**Fig. 10.**
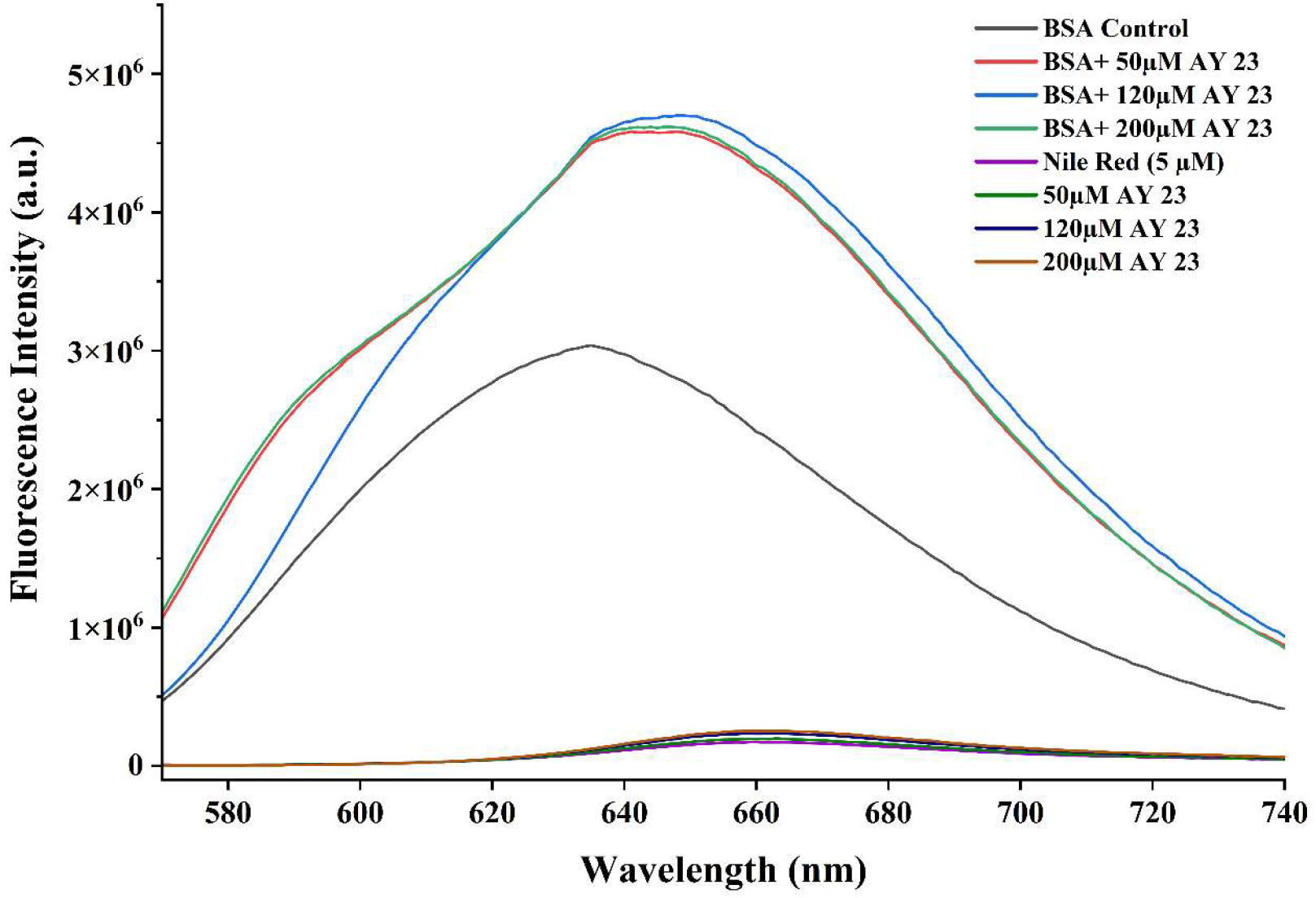
Nile red fluorescence spectra of BSA in the absence and presence of AY 23.

### 3.9. Thioflavin T Fluorescence Microscopy

Fluorescence microscopy was used to directly visualise the conformational changes and aggregation of BSA induced by AY 23, as shown in Fig. 11. The BSA control sample exhibited a dark, uniform field with no significant fluorescence, confirming the absence of aggregates in its native, soluble state. In stark contrast, the samples incubated with AY 23 showed a marked increase in fluorescence and distinct aggregates. The fluorescence signal is attributed to the AY 23 molecules incorporated within these structures. The images reveal the formation of fibrillar aggregates, which are elongated, thread-like structures characteristic of amyloid formation (He et al., 2022; Nirwal et al., 2021). This finding provides direct visual confirmation of the protein’s self-assembly, corroborating the results from the Nile red assay and DLS. The extent of fibril formation is directly correlated with the concentration of AY 23. The 50 µM AY 23 sample displayed a minimal number of small aggregates and short fibrils, suggesting the initial stages of aggregation. As the AY 23 concentration increased to 120 µM, a more extensive network of larger, denser fibrils became apparent. The sample with the highest concentration, 200 µM AY 23, exhibited the most extensive fibrillar network, with a high density of intertwined, interconnected fibrils, indicating a more advanced stage of aggregation. This suggests that AY 23 not only initiates but also accelerates the formation of these fibrillar structures in a dose-dependent manner. In addition to microscopy, ThT assay experiments were conducted, but because of the AY 23 broad absorption overlap (∼436 nm) with the ThT excitation range (Leulescu et al., 2018; Xue et al., 2017), we observed a typical ThT quenching pattern in BSA samples incubated with AY 23 (Supplementary Figure S3).

**Fig. 11.**
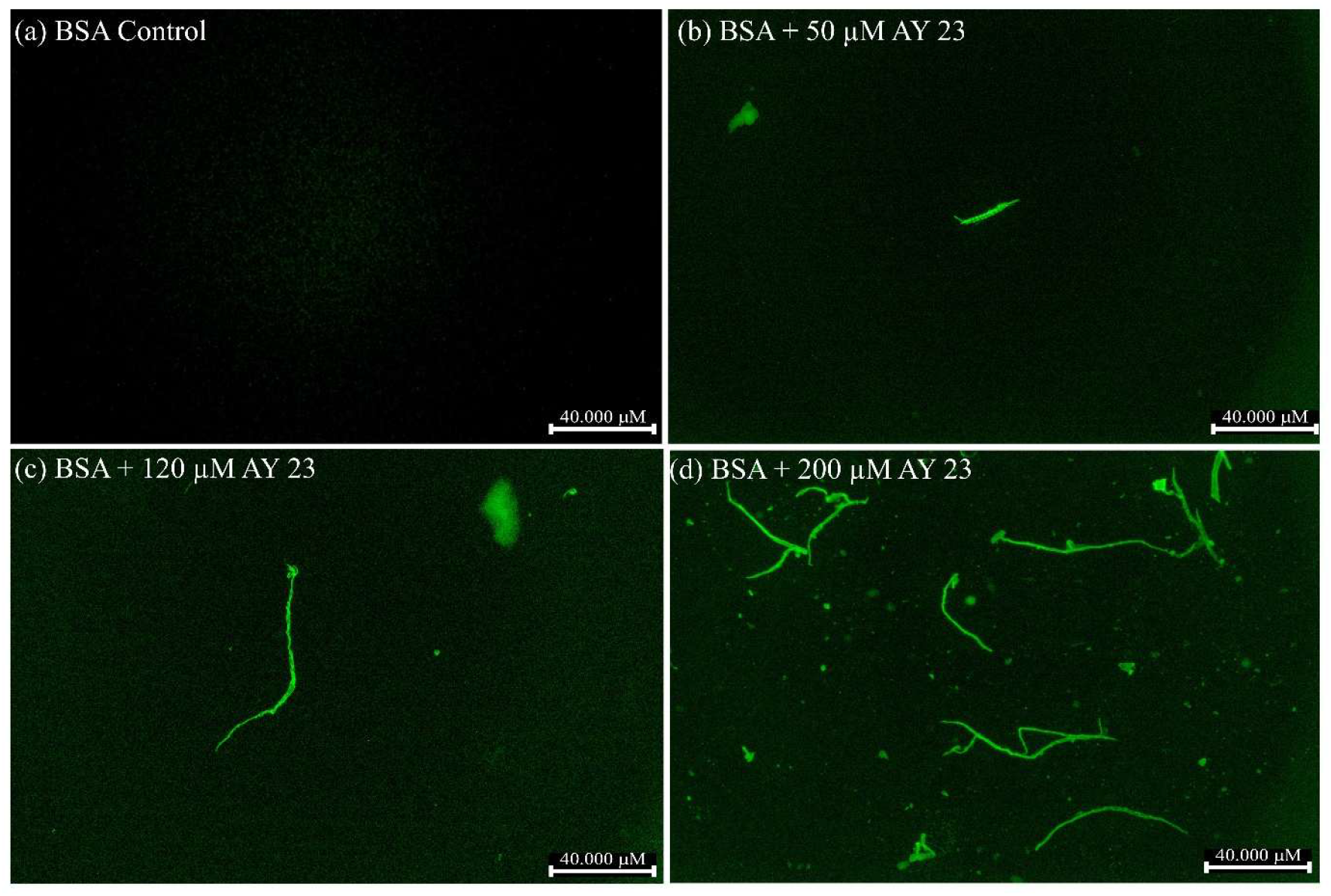
Fluorescence microscopy visualization of BSA conformational changes and aggregation induced by AY 23.

### 3.10. Atomic Force Microscopy (AFM)

Atomic Force Microscopy (AFM) was employed to visualise the morphological changes of BSA upon incubation with varying concentrations of AY 23, as shown in Fig. 12. The AFM image of the BSA control sample showed a homogeneous distribution of individual, globular particles with a uniform height, characteristic of monomeric BSA (Felix et al., 2026). This confirmed the protein’s native state in the absence of the dye. The addition of AY 23 resulted in a significant transition from a globular to an aggregated state. At the highest concentration of 200 µM AY 23, the AFM image revealed a profound transformation, with the entire field dominated by a dense and extensive network of elongated, rigid, and interconnected fibrils (Dubey et al., 2017). These observations provide direct visual evidence that AY 23 promotes the self-assembly of BSA into amyloid-like fibrillar aggregates, supporting findings from spectroscopic and light-scattering analyses.

**Fig. 12.**
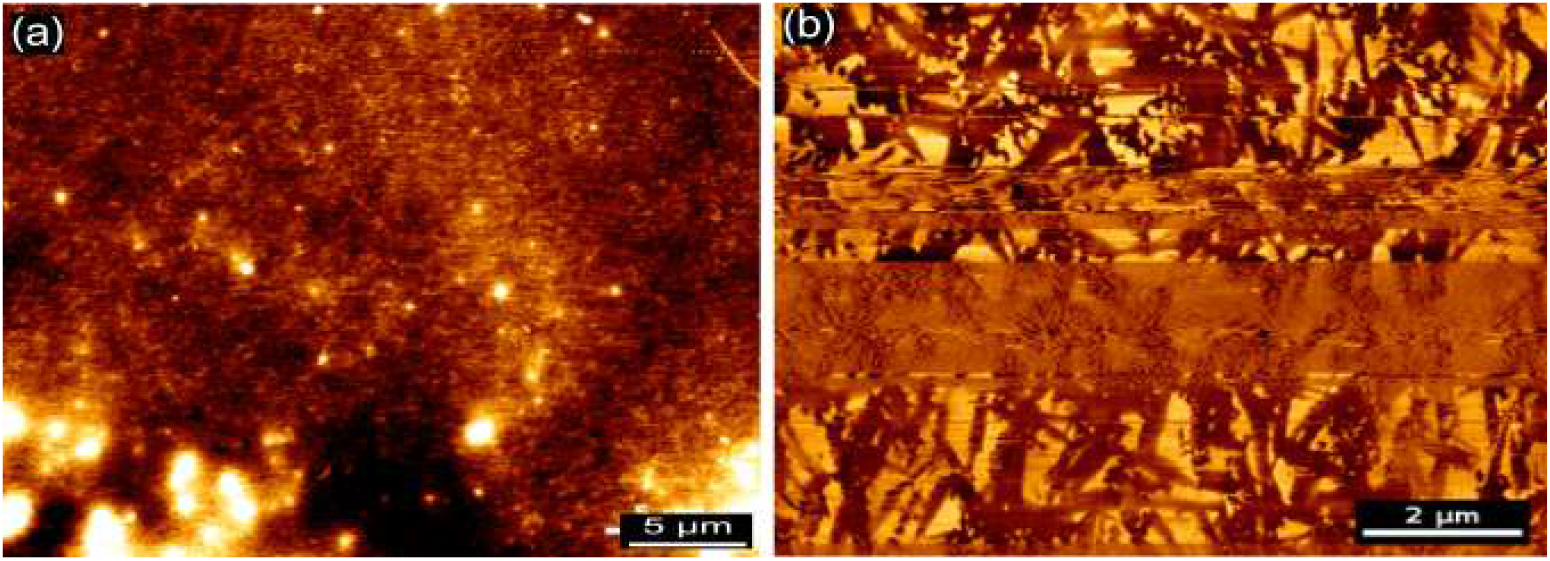
Atomic Force Microscopy (AFM) images showing morphological changes in BSA after incubation with varying concentrations of AY 23. (a) BSA Control and (b) BSA + 200 μM AY 23.

### 3.11. Hemolytic Assay

The erythrocytic potential of the AY 23-induced BSA fibrils was assessed by measuring the release of haemoglobin from human erythrocytes at an absorbance of 405 nm. The wavelength was selected due to its proximity to the high-intensity Soret band of haemoglobin, providing enhanced sensitivity for detecting subtle membrane disruptions (Greco et al., 2020). The experimental data depicted in Fig. 13 reveal a significant increase in hemolytic activity of AY 23-induced fibrils of BSA. The Fresh BSA remained in the nonhemolytic range at 1.19%. Also, we didn’t find hemolytic activity of independent AY 23 (data not shown). At day 9 of incubation, BSA-AY 23 samples showed 15.64% hemolysis at 50 µM and 33.73% at 120 µM. At the highest concentration of 200 µM, hemolysis was 42.26%, considerably higher compared to the lower concentrations of AY 23. These results indicate that the toxicity is largely driven by the extent of AY 23-induced fibrillation.

**Fig. 13.**
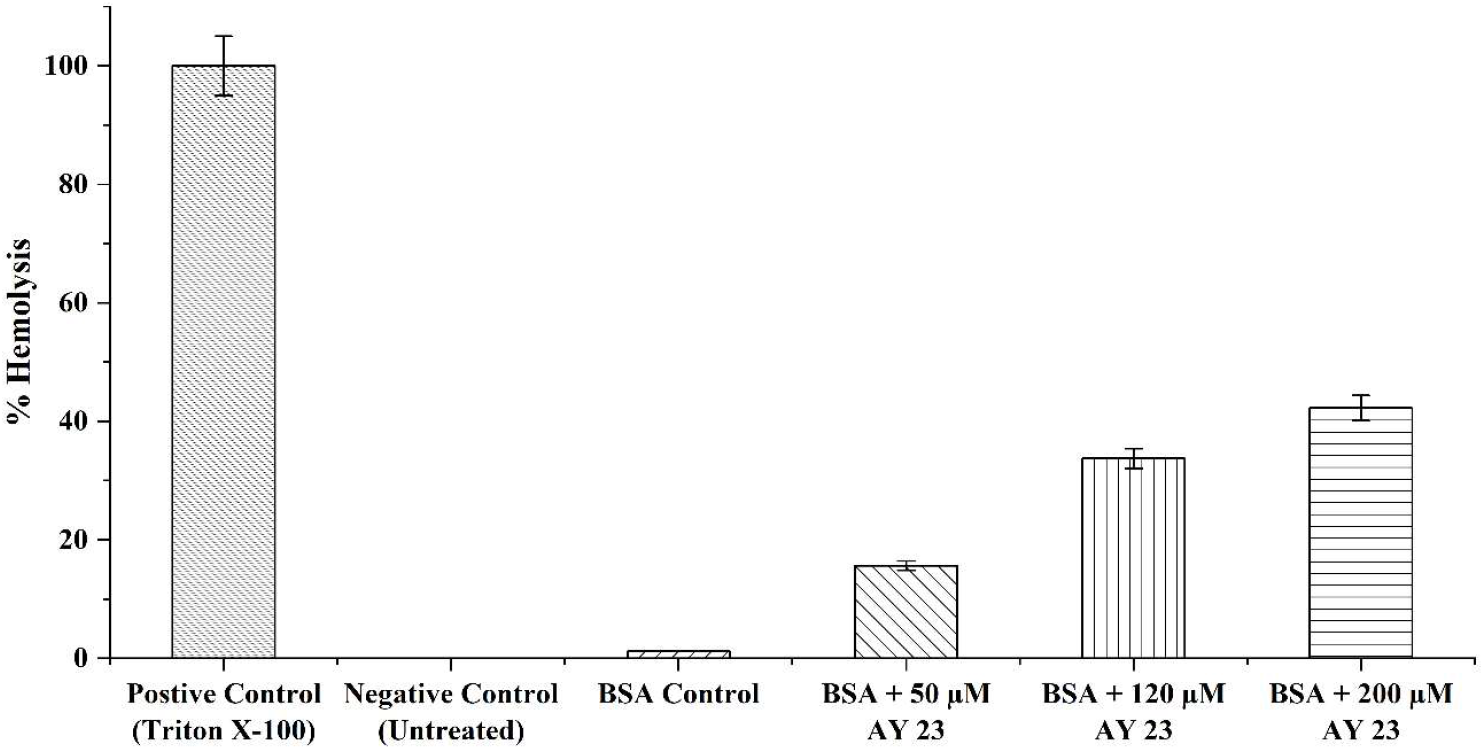
Hemolytic effect of AY 23 in the presence of BSA.

### 3.12. Proposed Mechanism of Interaction of Acid Yellow 23 (AY) and Bovine Serum Albumin (BSA)

The current study provides evidence of AY 23-mediated alterations in BSA that lead to the formation of toxic species. Tryptophan fluorescence quenching data for BSA with AY 23 suggest significant binding of the dye to the protein. This association is clarified by molecular docking simulations, which demonstrate that AY 23 binds spontaneously (ΔG = -9.7 kcal/mol) inside a hydrophobic pocket of BSA. Important interactions such as hydrogen bonds (e.g., GLU A:424, ASP A:111) and π-anion/hydrophobic forces involving aromatic rings and residues (e.g., ARG A:144) mediate this stable complex. The main cause of structural rearrangement is this strong molecular event, which gradually reduces the α-helical content (as shown by CD spectroscopy) and exposes hidden hydrophobic domains (as revealed by the Nile Red Assay). The main factor promoting self-assembly is this enhanced surface hydrophobicity, which causes misfolded monomers to clump together into sizable aggregates (Singh et al., 2021). The aggregation is quantitatively confirmed by the significant rise in hydrodynamic radius (DLS, up to ∼500 nm). Direct visualisation using fluorescence microscopy and atomic force microscopy provided compelling morphological confirmation of aggregation. The incorporation of AY 23 into these structures, as evidenced by fluorescence imaging, suggests that the dye acts as a structural modulator, stabilising the aggregated state. Finally, the hemolytic assay establishes the biological relevance of the AY 23-BSA interaction. The pronounced, concentration-dependent increase in hemolysis indicates that dye-induced aggregation significantly alters the protein’s biocompatibility. The transition to highly hemolytic behaviour at higher AY 23 concentrations suggests enhanced membrane-disruptive potential, likely arising from altered surface properties and the presence of rigid, multivalent aggregate assemblies. Therefore, the data support a mechanistic model in which AY 23 binds to BSA via hydrophobic and aromatic interactions, destabilises α-helices, promotes β-sheet-rich amyloid-like aggregation, and enhances cytotoxicity via fibrillation formed at higher concentrations. In contrast to the earlier serum albumin study (Al-Shabib et al., 2018a), which reported tatrazine (AY 23)-induced fibrillation only at pH 3.5, our results demonstrated AY 23’s capacity to cause molecular alterations and aggregation at pH 7.4 at micromolar concentrations. It is known that lower concentrations can lead to greater conformational selection, with longer interaction durations and high binding affinity (Deng et al., 2022; Greives and Zhou, 2014). These findings highlight the capacity of synthetic azo dyes to modulate protein structure and function, underscoring their potential toxicological relevance and the importance of evaluating dye–protein interactions in biological systems.

## 4. Conclusions

The work demonstrates that AY 23 substantially modifies the aggregation behaviour and structural integrity of BSA. By combining simulation, spectroscopic, microscopic, and biochemical methods, it was demonstrated that AY 23 forms stable complexes with BSA, induces the loss of the native α-helical structure, and promotes concentration- and time-dependent self-assembly into amyloid-like fibrillar aggregates enriched in β-sheets. The molecular docking analysis demonstrated that AY 23 binds favourably within a hydrophobic binding pocket of BSA, with a favourable predicted docking score of -9.7 kcal/mol, forming stabilising interactions with key residues including GLU424, ASP111, and ARG144, thereby supporting the strong protein–ligand association observed experimentally. The 100 ns MD simulation supported continued association of AY 23 with the predicted BSA binding region and identified recurring protein–ligand contacts during the trajectory. Significant alterations in surface hydrophobicity, aggregate architecture, and biological response were observed in tandem with these structural changes. Circular dichroism measurements, Nile red assay, and fluorescence quenching indicate that AY 23 binding induces significant conformational changes in BSA, followed by molecular alterations that lead to aggregate formation. SDS-PAGE analysis confirmed the presence of ordered aggregates induced by AY 23 interaction, while dynamic light scattering, fluorescence microscopy and atomic force microscopy confirmed the formation of large, heterogeneous aggregates. The biological ramifications of dye-induced protein aggregation are highlighted by the notable progressive increase in hemolytic activity observed with BSA fibrils at different AY 23 concentrations. This also indicates a significant decrease in protein biocompatibility at various dye levels. Overall, the results provide evidence for a mechanistic framework in which AY 23 perturbs the native structure of BSA and promotes amyloid-like self-assembly via hydrophobic and aromatic interactions, thereby serving as an inducer of protein aggregation. This study emphasises the need for a systematic assessment of dye–protein interactions under biologically relevant conditions, as well as the potential of synthetic azo dyes to affect protein structure and biological compatibility from health perspectives.

## Author Contributions

**Parshant Dahiya:** Methodology, Validation, Formal analysis, Investigation, Data curation, Writing - original draft, Software. **Alpana Verma:** Methodology, Validation, Formal analysis, Investigation. **Vishal Mevada:** Formal analysis, Visualisation**. Satish Kumar:** Formal analysis, Visualisation, Supervision. **Neelkant Verma:** Conceptualisation, Project administration, Visualisation, Funding acquisition, Supervision, Writing - review & editing, Methodology.

## Acknowledgements

The authors gratefully acknowledge the National Forensic Sciences University, Gandhinagar, Gujarat, for providing exceptional laboratory facilities, advanced instrumentation, an institutional fellowship, and a supportive research environment, which were instrumental in the successful completion of this study.

## Glossary

ADI: Acceptable Daily Intake
AFM: Atomic Force Microscopy
APS: Ammonium Persulfate
AU: Arbitrary Units
AY 23: Acid Yellow 23
BSA: Bovine Serum Albumin
CD: Circular Dichroism
DLS: Dynamic Light Scattering
Bis: N,N’-methylene bisacrylamide
PDB: Protein Data Bank
PDI: Polydispersity Index
Rh: Hydrodynamic Radii
SDS: Sodium Dodecyl Sulfate
SDS-PAGE: Sodium Dodecyl Sulfate–Polyacrylamide Gel Electrophoresis
TEMED: N,N,N’, N’-tetramethylethylenediamine
ThT: Thioflavin T
Trp: Tryptophan UV Ultraviolet
RMSD: Root Mean Square Deviation
RMSF: Root Mean Square Fluctuation

## References

Abd-Elhakim, Y.M., Hashem, M.M., El-Metwally, A.E., Anwar, A., Abo-EL-Sooud, K., Moustafa, G.G., Ali, H.A., 2018. Comparative haemato-immunotoxic impacts of long-term exposure to tartrazine and chlorophyll in rats. International Immunopharmacology 63, 145–154. 10.1016/j.intimp.2018.08.002.

Al-Shabib, N.A., Khan, J.M., Alsenaidy, M.A., Alsenaidy, A.M., Khan, M.S., Husain, F.M., Khan, M.R., Naseem, M., Sen, P., Alam, P., Khan, R.H., 2018a. Unveiling the stimulatory effects of tartrazine on human and bovine serum albumin fibrillogenesis: Spectroscopic and microscopic study. Spectrochimica acta. Part A, Molecular and biomolecular spectroscopy 191, 116–124. 10.1016/j.saa.2017.09.062.

Al-Shabib, N.A., Khan, J.M., Malik, A., Alsenaidy, M.A., Rehman, M.T., AlAjmi, M.F., Alsenaidy, A.M., Husain, F.M., Khan, R.H., 2018b. Molecular insight into binding behavior of polyphenol (rutin) with beta lactoglobulin: Spectroscopic, molecular docking and MD simulation studies. Journal of Molecular Liquids 269, 511–520. 10.1016/j.molliq.2018.07.122.

Amchova, P., Kotolova, H., Ruda-Kucerova, J., 2015. Health safety issues of synthetic food colorants. Regulatory toxicology and pharmacology : RTP 73, 914–922. 10.1016/j.yrtph.2015.09.026.

Amchova, P., Siska, F., Ruda-Kucerova, J., 2024. Safety of tartrazine in the food industry and potential protective factors. Heliyon 10, e38111. 10.1016/j.heliyon.2024.e38111.

Ansari, N.K., Hasan, S., Siddiqui, G.A., Naeem, A., 2025. Protective Effect of Curcumin on Thermally Aggregated Bovine Serum Albumin. Cell Biochem Biophys 83, 4897–4906. 10.1007/s12013-025-01810-6.

Axon, A., May, F.E.B., Gaughan, L.E., Williams, F.M., Blain, P.G., Wright, M.C., 2012. Tartrazine and sunset yellow are xenoestrogens in a new screening assay to identify modulators of human oestrogen receptor transcriptional activity. Toxicology 298, 40–51. 10.1016/j.tox.2012.04.014.

Benkhaya, S., M’rabet, S., El Harfi, A., 2020. Classifications, properties, recent synthesis and applications of azo dyes. Heliyon 6, e03271. 10.1016/j.heliyon.2020.e03271.

Branen, A., Davidson, P., Salminen, S., Thorngate III, J., 2001. Food Additives. CRC Press.

Cao, H.-W., Chen, Y.-S., Li, J.-Z., Chen, H.-W., Li, L.-Y., Li, Z.-K., Wang, M.-Q., 2024. Development of D-π-A organic dyes for discriminating HSA from BSA and study on dye-HSA interaction. Bioorganic Chemistry 147, 107360. 10.1016/j.bioorg.2024.107360.

Carter, D.C., Ho, J.X., 1994. Structure of Serum Albumin, in: Anfinsen, C.B., Edsall, J.T., Richards, F.M., Eisenberg, D.S. (Eds.), Advances in Protein Chemistry : Lipoproteins, Apolipoproteins, and Lipases, vol. 45. Academic Press, pp. 153–203.

Castro, G.R., Larson, B.K., Panilaitis, B., Kaplan, D.L., 2005. Emulsan quantitation by Nile red quenching fluorescence assay. Appl Microbiol Biotechnol 67, 767–770. 10.1007/s00253-004-1849-2.

Demeule, B., Gurny, R., Arvinte, T., 2007. Detection and characterization of protein aggregates by fluorescence microscopy. International Journal of Pharmaceutics 329, 37–45. 10.1016/j.ijpharm.2006.08.024.

Deng, Y., Efremov, A.K., Yan, J., 2022. Modulating binding affinity, specificity, and configurations by multivalent interactions. Biophysical Journal 121, 1868–1880. 10.1016/j.bpj.2022.04.017.

Dezhampanah, H., Mohammadi, A., Mousazadeh Moghaddam Pour, A., 2023. Investigation on intermolecular interaction of synthesized azo dyes with bovine serum albumin. Journal of biomolecular structure & dynamics 41, 970–981. 10.1080/07391102.2021.2015444.

Dubey, K., Anand, B.G., Shekhawat, D.S., Kar, K., 2017. Eugenol prevents amyloid formation of proteins and inhibits amyloid-induced hemolysis. Sci Rep 7, 40744. 10.1038/srep40744.

El-sakhawy, M.A., Mohamed, D.W., Ahmed, Y.H., 2019. Histological and immunohistochemical evaluation of the effect of tartrazine on the cerebellum, submandibular glands, and kidneys of adult male albino rats. Environ Sci Pollut Res 26, 9574–9584. 10.1007/s11356-019-04399-5.

Felix, M., Lendel, C., Fielden, M., Carrera-Sanchez, C., Rutland, M., 2026. Reinforcement of bovine serum albumin-based interfaces by the addition of genipin and carboxymethyl cellulose: Effect of pH. Journal of Colloid and Interface Science 723, 140909. 10.1016/j.jcis.2026.140909.

Greco, I., Molchanova, N., Holmedal, E., Jenssen, H., Hummel, B.D., Watts, J.L., Håkansson, J., Hansen, P.R., Svenson, J., 2020. Correlation between hemolytic activity, cytotoxicity and systemic in vivo toxicity of synthetic antimicrobial peptides. Sci Rep 10, 13206. 10.1038/s41598-020-69995-9.

Greives, N., Zhou, H.-X., 2014. Both protein dynamics and ligand concentration can shift the binding mechanism between conformational selection and induced fit. Proceedings of the National Academy of Sciences of the United States of America 111, 10197–10202. 10.1073/pnas.1407545111.

Halder, S., Aggrawal, R., Jana, S., Saha, S.K., 2021. Binding interactions of cationic gemini surfactants with gold nanoparticles-conjugated bovine serum albumin: A FRET/NSET, spectroscopic, and docking study. Journal of photochemistry and photobiology. B, Biology 225, 112351. 10.1016/j.jphotobiol.2021.112351.

He, L., Liu, Y., Xu, J., Li, J., Cheng, G., Cai, J., Dang, J., Yu, M., Wang, W., Duan, W., Liu, K., 2022. Inhibitory Effects of Myriocin on Non-Enzymatic Glycation of Bovine Serum Albumin. Molecules 27, 6995. 10.3390/molecules27206995.

Kawachi, Y., Kameyama, R., Handa, A., Takahashi, N., Tanaka, N., 2013. Role of the N-terminal amphiphilic region of ovalbumin during heat-induced aggregation and gelation. J. Agric. Food Chem. 61, 8668–8675. 10.1021/jf402456v.

Keener, L., 2022. Chapter 14 - Food safety and regulatory survey of food additives and other substances in human food, in: Martinović, A., Oh, S., Lelieveld, H. (Eds.), Ensuring Global Food Safety (Second Edition). Academic Press, pp. 259–273.

König, J., 2015. 2 - Food colour additives of synthetic origin, in: Scotter, M.J. (Ed.), Colour Additives for Foods and Beverages : Woodhead Publishing Series in Food Science, Technology and Nutrition. Woodhead Publishing, Oxford, pp. 35–60.

Koundal, S., Pathania, A., Kour, H.D., Pathania, A.R., Kaur, J., Juneja, B., Sharma, G.C., Bhowmik, A., Santhosh, A.J., 2025. Molecular interaction study of L-Ornithine with bovine serum albumin using spectroscopic and molecular docking methods. Sci Rep 15, 11997. 10.1038/s41598-025-93108-z.

Lakowicz, J.R., 2006. Principles of fluorescence spectroscopy, 3rd ed. Springer, New York, xxvi, 954.

Leulescu, M., Rotaru, A., Pălărie, I., Moanţă, A., Cioateră, N., Popescu, M., Morîntale, E., Bubulică, M.V., Florian, G., Hărăbor, A., Rotaru, P., 2018. Tartrazine: physical, thermal and biophysical properties of the most widely employed synthetic yellow food-colouring azo dye. J Therm Anal Calorim 134, 209–231. 10.1007/s10973-018-7663-3.

Li, X., Fang, Y., Al-Assaf, S., Phillips, G.O., Yao, X., Zhang, Y., Zhao, M., Zhang, K., Jiang, F., 2012. Complexation of bovine serum albumin and sugar beet pectin: structural transitions and phase diagram. Langmuir 28, 10164–10176. 10.1021/la302063u.

Liu, Y., Gan, N., Liu, P., Xu, J., Geng, F., Di Wu, 2026. Characterization of the interaction between tartrazine and β-lactoglobulin under different pH conditions: multispectral analysis and computer simulation. Current Research in Food Science 12, 101323. 10.1016/j.crfs.2026.101323.

Mavani, A., Ray, D., Aswal, V.K., Bhattacharyya, J., 2023. Understanding the molecular interaction of BSA protein with antibiotic sulfa molecule(s) for novel drug development. Journal of Molecular Structure 1287, 135697. 10.1016/j.molstruc.2023.135697.

Micsonai, A., Wien, F., Murvai, N., Nyiri, M.P., Balatoni, B., Lee, Y.-H., Molnár, T., Goto, Y., Jamme, F., Kardos, J., 2025. BeStSel: analysis site for protein CD spectra-2025 update. Nucleic Acids Res 53, W73–W83. 10.1093/nar/gkaf378.

Mota, I.G.C., Neves, R.A.M.D., Da Nascimento, S.S.C., Maciel, B.L.L., Morais, A.H.D.A., Passos, T.S., 2023. Artificial Dyes: Health Risks and the Need for Revision of International Regulations. Food Reviews International 39, 1578–1593. 10.1080/87559129.2021.1934694.

Nirwal, S., Bharathi, V., Patel, B.K., 2021. Amyloid-like aggregation of bovine serum albumin at physiological temperature induced by cross-seeding effect of HEWL amyloid aggregates. Biophysical Chemistry 278, 106678. 10.1016/j.bpc.2021.106678.

Pan, X., Qin, P., Liu, R., Wang, J., 2011. Characterizing the Interaction between tartrazine and two serum albumins by a hybrid spectroscopic approach. J. Agric. Food Chem. 59, 6650–6656. 10.1021/jf200907x.

Pignataro, M.F., Herrera, M.G., Dodero, V.I., 2020. Evaluation of Peptide/Protein Self-Assembly and Aggregation by Spectroscopic Methods. Molecules 25, 4854. 10.3390/molecules25204854.

Ramesh, H., Bhuyan, A.K., 2024. Food Colorants Acid Yellow 23, Azorubine, and Acid Red 18 Cause Aggregation and Fibrillation of Proteins. ACS Food Sci. Technol. 4, 963–974. 10.1021/acsfoodscitech.3c00707.

Roy, A., Tiwari, S., Karmakar, S., Anki Reddy, K., Pandey, L.M., 2019. The effect of the stoichiometric ratio of zinc towards the fibrillation of Bovine Serum Albumin (BSA): A mechanistic insight. International Journal of Biological Macromolecules 123, 409–419. 10.1016/j.ijbiomac.2018.11.120.

Sackett, D.L., Wolff, J., 1987. Nile red as a polarity-sensitive fluorescent probe of hydrophobic protein surfaces. Analytical Biochemistry 167, 228–234. 10.1016/0003-2697(87)90157-6.

Sæbø, I.P., Bjørås, M., Franzyk, H., Helgesen, E., Booth, J.A., 2023. Optimization of the Hemolysis Assay for the Assessment of Cytotoxicity. International Journal of Molecular Sciences 24, 2914. 10.3390/ijms24032914.

Singh, G., Kaur, M., Singh, M., Kaur, H., Kang, T.S., 2021. Spontaneous Fibrillation of Bovine Serum Albumin at Physiological Temperatures Promoted by Hydrolysis-Prone Ionic Liquids. Langmuir 37, 10319–10329. 10.1021/acs.langmuir.1c01350.

Varma, N., Singh, I., Dahiya, M.S., Ravi, V.K., Kumar, S., 2018. Structural perturbation by arsenic triggers the aggregation of hen egg white lysozyme by promoting oligomers formation. International Journal of Biological Macromolecules 109, 1108–1114. 10.1016/j.ijbiomac.2017.11.096.

Waghmare, M., Khade, B., Chaudhari, P., Dongre, P., 2018. Multiple layer formation of bovine serum albumin on silver nanoparticles revealed by dynamic light scattering and spectroscopic techniques. J Nanopart Res 20, 185. 10.1007/s11051-018-4286-3.

Wang, W., 2005. Protein aggregation and its inhibition in biopharmaceutics. International Journal of Pharmaceutics 289, 1–30. 10.1016/j.ijpharm.2004.11.014.

Xue, C., Lin, T.Y., Chang, D., Guo, Z., 2017. Thioflavin T as an amyloid dye: fibril quantification, optimal concentration and effect on aggregation. R. Soc. Open Sci. 4, 160696. 10.1098/rsos.160696.

