## Supplementry Data for "Molecular Alterations of Bovine Serum Albumin Induced by the Food Dye Acid Yellow 23: A Mechanistic Study"

**Supplementary material**

**Supplementary Table 1**. The estimated secondary structure content (%) of BSA + AY 23 (Various concentrations) from the obtained CD spectra using the BeStSel CD spectrum analysis tool.

| **Sample** | **Helix** | **Antiparallel** | **Parallel** | **Turn** | **Others** |
| --- | --- | --- | --- | --- | --- |
| BSA Control | 58.7 | 0 | 0 | 8.1 | 33.2 |
| BSA + 50μM AY 23 | 36.1 | 3.9 | 22.9 | 1.3 | 35.9 |
| BSA + 120μM AY 23 | 33.7 | 5.7 | 21.7 | 4.1 | 34.8 |
| BSA + 200μM AY 23 | 19.5 | 4.2 | 5.3 | 11.5 | 59.4 |

**Supplementary Figure 1**. UV–Vis Absorption Spectra of BSA (30 µM), AY 23 alone (30 µM), BSA-AY 23 complex (1:1) and difference Spectra of BSA-AY 23.

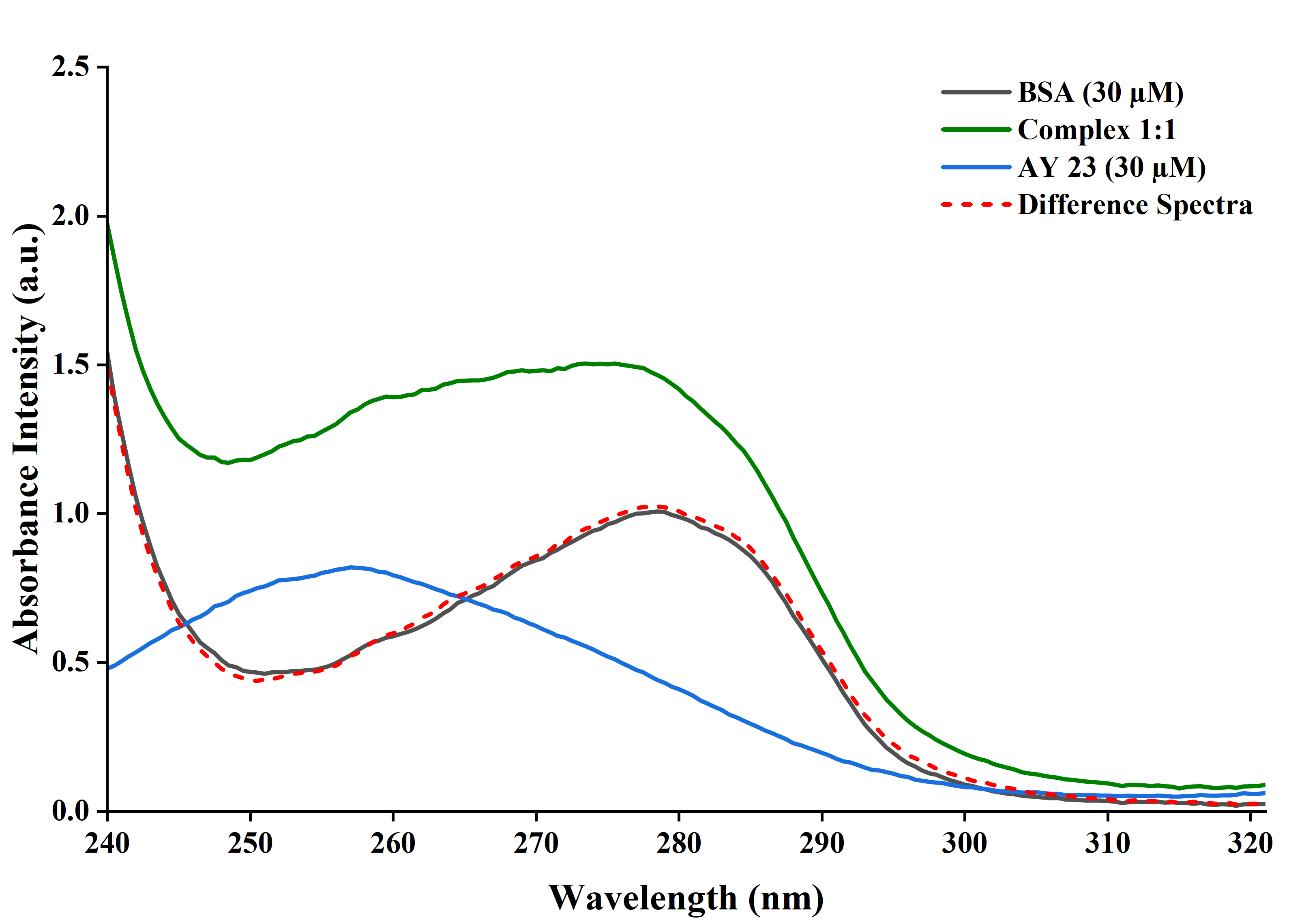

**Supplementary Figure 2:** Intrinsic tryptophan fluorescence emission spectra of BSA in the presence and absence of Acid Yellow 23 (10–210 µM.

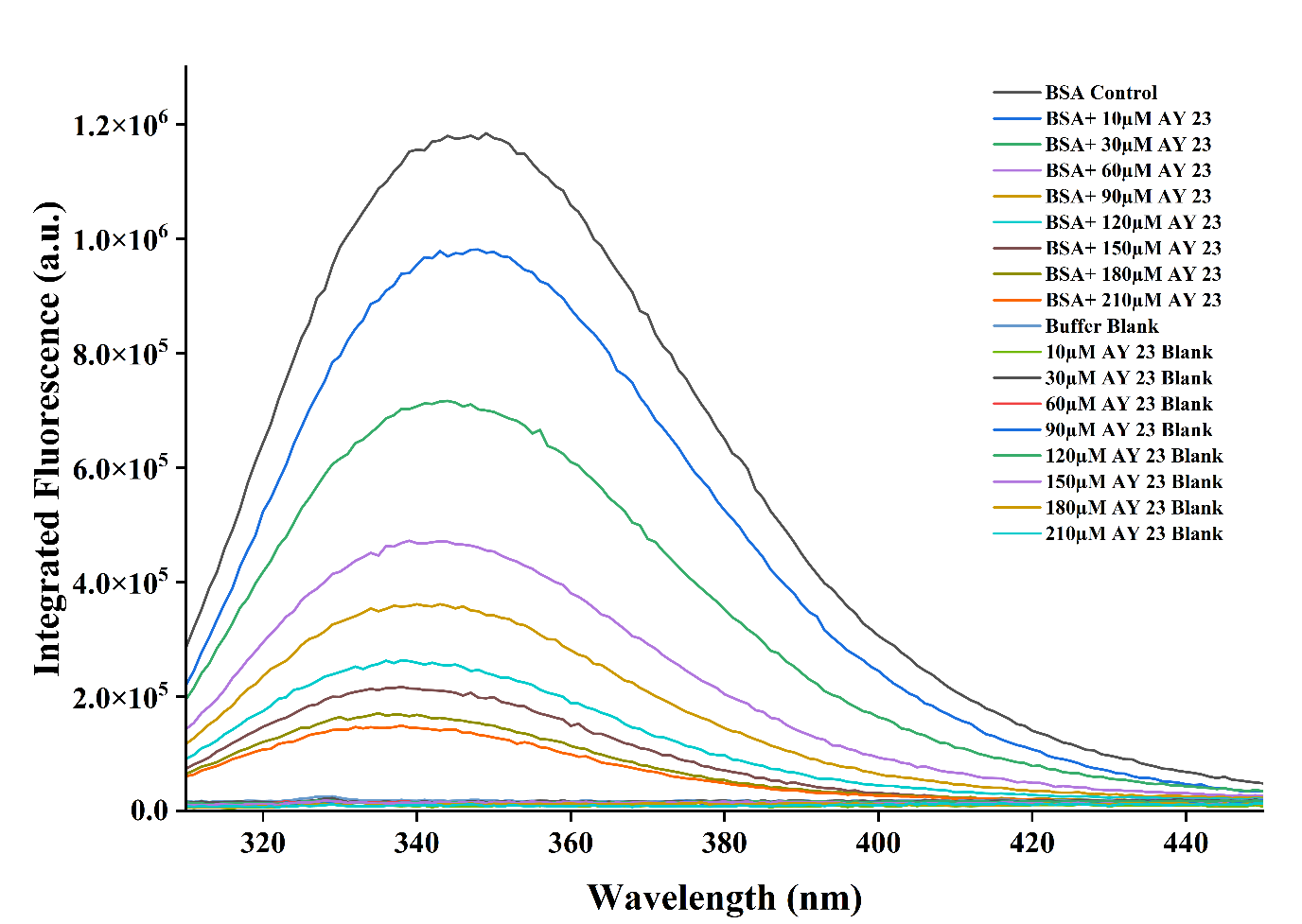

**Supplementary Figure 3:** ThT fluorescence intensity of BSA with and without Acid Yellow 23 (50, 120, and 200 µM) over the period of 9 days.

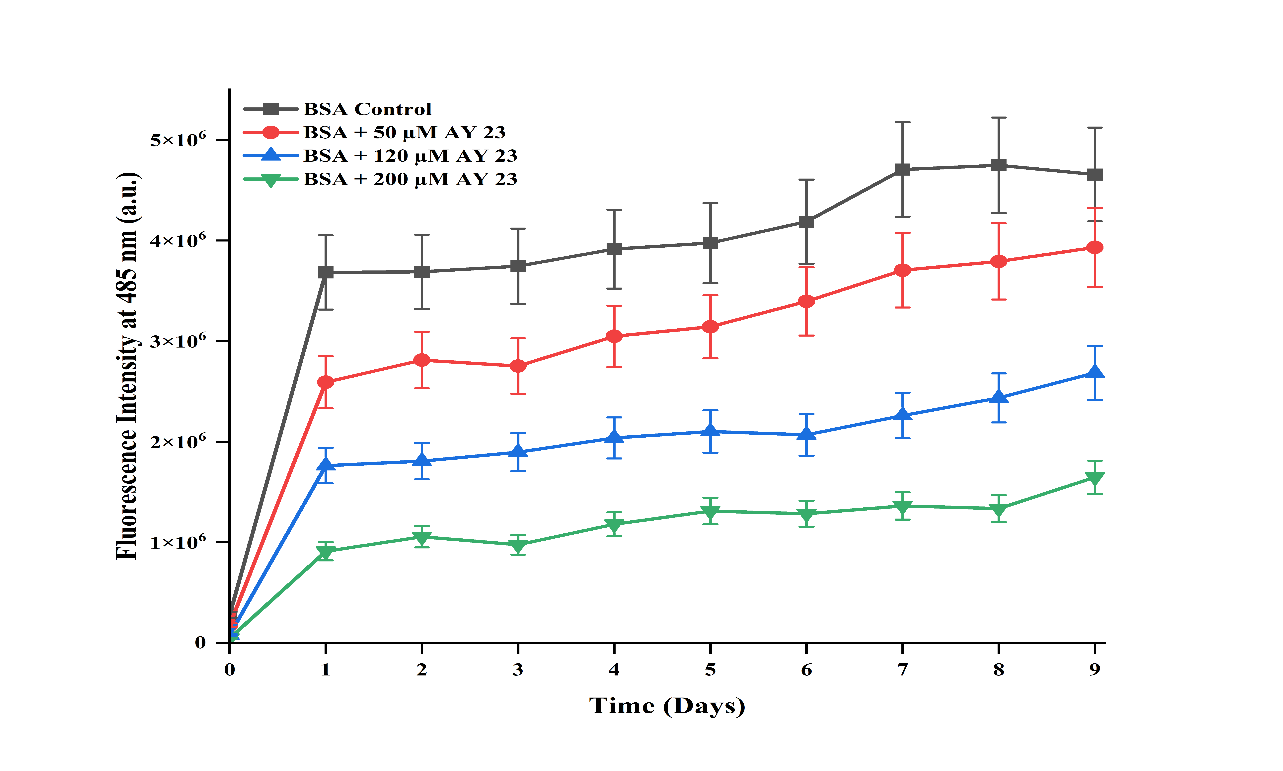
